# A Plant Flavonol Rescues a Pathogenic Mutation Associated with Kinesin in Neurons

**DOI:** 10.1101/2023.07.19.548188

**Authors:** Yongping Chai, Dong Li, Weibin Gong, Jingyi Ke, Dianzhe Tian, Zhe Chen, Angel Guo, Zhengyang Guo, Wei Li, Wei Feng, Guangshuo Ou

## Abstract

KIF1A, a microtubule-based motor protein responsible for axonal transport, is linked to a group of neurological disorders known as KIF1A-associated neurological disorder (KAND). Current therapeutic options for KAND are limited. Here, we introduced the clinically relevant KIF1A(R11Q) variant into the *C. elegans* homolog UNC-104, resulting in uncoordinated animal behaviors. Through genetic suppressor screens, we identified intragenic mutations in UNC-104’s motor domain that rescued synaptic vesicle localization and coordinated movement. We showed that two suppressor mutations partially recovered motor activity in vitro by counteracting the structural defect caused by R11Q at KIF1A’s nucleotide-binding pocket. We found that supplementation with fisetin, a plant flavonol, improved KIF1A(R11Q) worms’ movement and morphology. Notably, our biochemical and single-molecule assays revealed that fisetin directly restored the ATPase activity and processive movement of human KIF1A(R11Q) without affecting wild-type KIF1A. These findings suggest fisetin as a potential intervention for enhancing KIF1A(R11Q) activity and alleviating associated defects in KAND.

## Introduction

Axonal transport, facilitated by microtubules (MTs), is essential for neural development and function^1–3^. It involves bidirectional movement mediated by kinesin superfamily proteins (KIFs) and cytoplasmic dynein-1^4–6^. Kinesin-3 family motors, such as UNC-104, transport synaptic vesicles (SVs) from the cell body to the axon^7, 8^. Mutations in UNC-104 result in abnormal SV localization and accumulation^9, 10^. UNC-104/KIF1A consists of various domains, including a motor domain, neck coil domain (NC), coiled-coil domains, forkhead-associated (FHA) domains, and a C-terminal pleckstrin homology (PH) domain^11–13^. Genetic links have been established between KIF1A mutations and impaired axonal transport, leading to neurodegeneration and KIF1A-associated neurological disorders (KAND)^14–19^. Numerous KIF1A mutations have been identified in patients, associated with conditions ranging from congenital neuropathies to hereditary spastic paraplegia (SPG)^16–18, 20–23^. However, effective interventions for these mutations and KAND therapies are still largely unknown.

*C. elegans* has emerged as a valuable genetic model for studying KAND. The functional defects observed in the *unc-104* mutant were rescued by introducing the human homolog KIF1A into *C. elegans*, indicating the evolutionary conservation of these motor proteins^24^. Recent research has utilized genome editing techniques to introduce KAND-associated mutations into the *C. elegans* genome, allowing for the systematic analysis of axonal transport and animal behavior^25, 26^. Complemented by biochemical and single-molecule assays conducted on KIF1A *in vitro*, these studies shed light on the dominant nature of *de novo* KAND variants and reveal that certain mutations lead to an abnormal increase in KIF1A motor activity^25, 26^. This suggests that hyperactivation of the motor, rather than loss-of-function, may contribute to motor neuron diseases.

*C. elegans* serves as an excellent platform not only for modeling KAND mutations but also for conducting genetic suppressor screens, offering valuable insights into potential interventions for KAND. For instance, one screen uncovered UNC-16, a kinesin cargo adaptor protein, and specific amino acid substitutions in the motor stalk region as intergenic suppressors capable of restoring the loss-of-function phenotype of *unc-104 (e1265)*^27, 28^. Another unbiased screen identified a gain-of-function allele of *unc-104* as a suppressor that restored presynaptic patterning and axonal transport disrupted by the loss of ARL-8, an Arf-like small G protein^29^. These findings demonstrate the feasibility of isolating suppressors for *unc-104* from both loss- and gain-of-function contexts. Additionally, a recent screen successfully identified a mutation that partially restored the motility of a clinical KIF1A variant with reduced motor activity^26^.

This study investigates the impact of the R11Q mutation in the KIF1A motor domain, which has been observed in a patient with combined Autism Spectrum Disorder (ASD) and attention-deficit/hyperactivity disorder^16^. The mutation disrupts the stability of the nucleotide-binding pocket, potentially affecting ATP binding or ADP release^13^. The recovery mechanism for such a detrimental mutation in the ATP/ADP-binding pocket remains unclear. Through genetic suppressor screens, biochemical assays, and single-molecule assays, we demonstrate that introducing additional mutations in the nucleotide-binding pocket restores the defects caused by R11Q, likely by restoring the pocket’s function. While other substitutions in the motor domain do not restore motility in vitro, they do rescue synaptic and coordination defects in the animal model. Furthermore, treatment with the plant flavonol fisetin improves movement in R11Q worms as well as the *in vitro* motor activity of the human KIF1A(R11Q) without affecting the wild-type KIF1A, which suggests a potential therapeutic approach for KAND. Our findings shed light on multiple strategies to address R11Q defects at both biochemical and animal levels, offering insights into KAND therapies.

## Results

### *C. elegans* genetic screens identified suppressors for the KIF1A(R11Q) mutation

To investigate the molecular factors involved in the cellular deficits caused by the disease-associated mutation KIF1A(R11Q), we employed the CRISPR-Cas9 genome editing technique to generate a *C. elegans* model. We introduced the corresponding *unc-104(R9Q)* mutation into the UNC-104::GFP knock-in strain (Fig. 1A-C). The resulting *unc-104(R9Q)* mutant animals displayed severe coiling, shortened body length, and pronounced impairments in coordinated movement, with a 100% occurrence rate (Fig. 1D, n > 200). These observed phenotypes closely resembled those exhibited by previously reported loss-of-function mutations identified in earlier studies ^26^.

**Fig.1.**
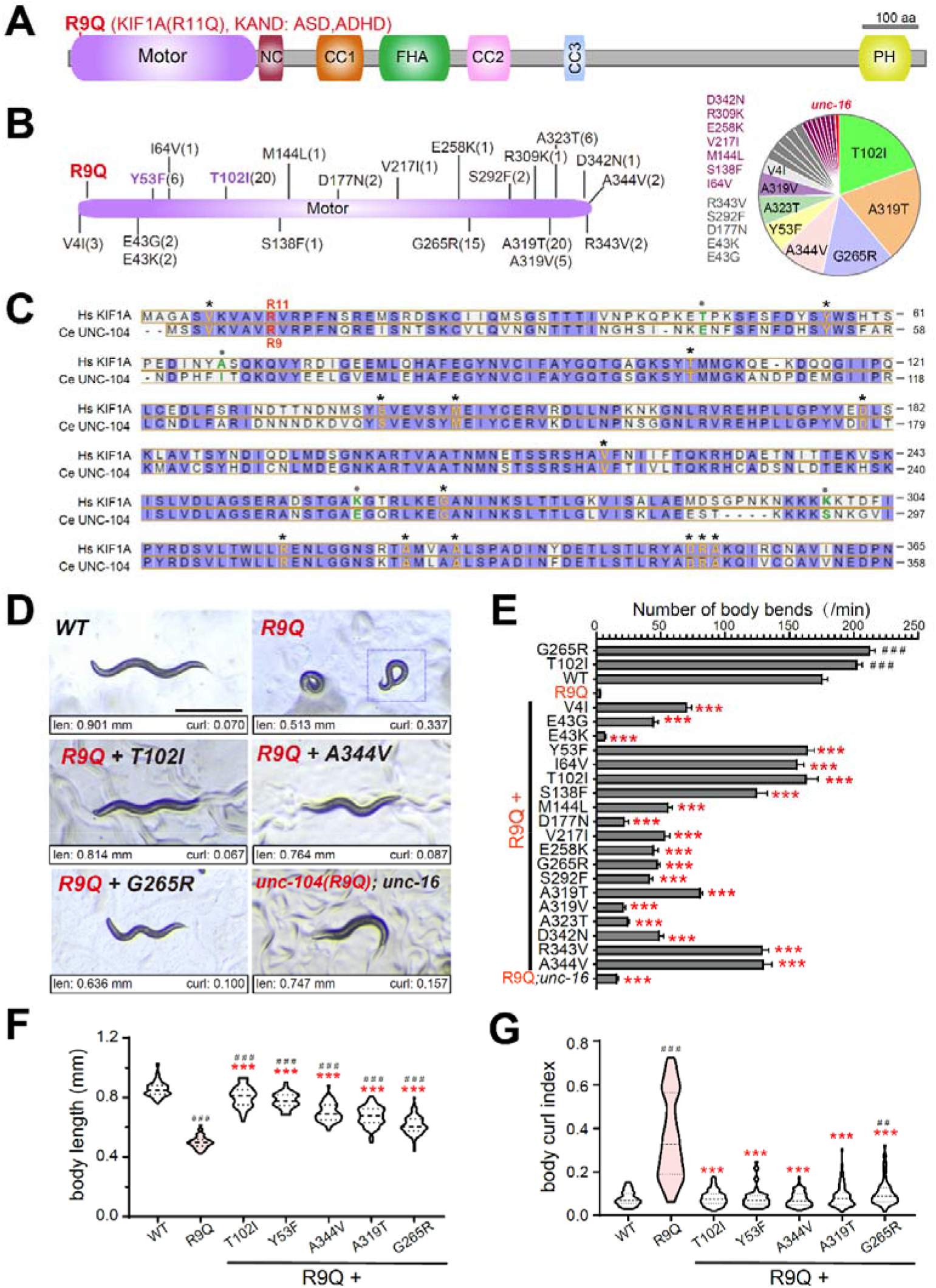
Genetic suppressor screens for UNC-104(R9Q) (A) Schematic of the domain organization of UNC-104 motor protein. NC, neck coiled-coil. CC, coiled-coil. FHA, Forkhead-associated. PH, pleckstrin homology. The R9Q mutation, which corresponds to the KIF1A-associated neuronal disorder (KAND)-associated KIF1A mutation R11Q, was introduced into the UNC-104 protein using the CRISPR-Cas9-based genome editing method. (B) Left: Amino acid location of isolated R9Q intragenic suppressors within the UNC-104 motor domain. The number of suppressors carrying each mutation site is given in parentheses. Right: pie chart exhibiting the relative population of individual variants among all suppressors. (C) Alignment of motor domain sequence between human KIF1A and *C. elegans* UNC-104 proteins. Identical residues are highlighted in purple, and intragenic suppressor sites are labeled with an asterisk or filled circle if they are identical or similar between KIF1A and UNC-104. (D) Representative bright field images of WT, *unc-104(R9Q)* mutant, and homozygous suppressor animals at the L4 larval stage. The measured body length (len) and calculated curl index (curl) are shown at the bottom of each image. The data shown in the R9Q image were obtained from the animal framed. Scale bar, 0.5 mm. Fig. S1 shows the definition of the curl index and more representative images. (E) The number of animal body bends in a water drop was counted for one minute and shown in columns. Identified UNC-104 amino acid changes in the examined animals are indicated. Error bars represent the SEM, n = 30 animals. Statistical significance was calculated by the unpaired Student’s *t*-test: *** *p* < 0.001 compared to the animal carrying UNC-104(R9Q) mutation, ^#^ ^#^ ^#^ *p* < 0.001 compared to the WT animals. (F-G)Quantification of animal body length (F) and curl index (G) of five representative suppressors that partially rescued morphological phenotypes of *unc-104(R9Q)* animals. *** *p* < 0.001, unpaired Student’s *t*-test compared with *unc-104(R9Q)* mutant animals; ^#^ ^#^ *p* < 0.01, ^#^ ^#^ ^#^ *p* < 0.001, unpaired Student’s *t*-test compared with WT animals.

The complete penetrance of animal defects in the *unc-104(R9Q)* mutant enabled us to conduct a large-scale and high-throughput suppressor screen. However, the *unc-104(R9Q)* mutant animals exhibited limited mobility and had a reduced brood size compared to wild-type (WT) N2 animals (56.2 ± 23.8, mean ± SD, n=18, approximately one-quarter of the brood size of wild-type animals), which presented challenges in obtaining sufficient synchronized progenies for chemical mutagenesis using conventional methods. To overcome this hurdle, we developed a specialized filter apparatus with a 15 μm diameter that allowed for the enrichment of L1 larvae from mixed population cultures based on their sizes (see Materials and Methods). Through this approach, we isolated a total of 103 independent suppressor mutants of *unc-104(R9Q)* from approximately 100,000 mutagenized haploid genomes. Whole-genome sequencing and bioinformatics analyses revealed 20 intragenic missense mutations within the motor domain, with most of the altered residues highly conserved between *C. elegans* and humans (Fig. 1B-C). Additionally, we identified an intergenic suppressor allele, *unc-16(E76K)*, which encodes a conserved JNK-signaling scaffold protein (Fig. 1B). Although the rescue efficiency of the intergenic suppressor was weaker compared to the intragenic suppressors (Fig. 1D-E), it still demonstrated significant effects. Notably, a previously reported missense mutation, L75P, in *unc-16* was also found to be a suppressor of other *unc-104* partial loss-of-function alleles^27^.

Through our genetic screens, we identified multiple alleles carrying the same intragenic mutations that exhibited identical rescue effects (Fig. 1B), suggesting that highly recurrent mutations may be responsible for recovering the UNC phenotypes in UNC-104(R9Q). To experimentally validate these identified mutations, we introduced *unc-104(T102I)* or *unc-104(G265R)* lesions into wild-type animals using genome editing. Neither of these mutant strains displayed any noticeable movement or morphology defects (Fig. 1E). We then introduced the *unc-104(R9Q)* mutation into these two strains, and interestingly, the resulting double mutant animals did not exhibit UNC defects (Fig. S1A). This phenocopied the observations made with the *unc-104(R9QT102I)* and *unc-104(R9QG265R)* mutants obtained from our genetic suppressor screens. These results confirm that the T102I and G265R amino acid substitutions are indeed intragenic mutations that restore the animal defects caused by the R9Q mutation in UNC-104.

To assess the recovery of worm motility, we conducted an animal swimming assay ^25, 26, 30^ and compared the paralyzed *unc-104(R9Q)* animals with the suppressor strains. All suppressors showed significant improvements in animal motility, although the levels of improvement varied (Fig. 1E). Among the five selected suppressors (Y53F, T102I, A344V, A319T, and G265R), we evaluated their effects on body length and animal morphology (Fig. 1F-G and Fig. S1B-C). The body length of *unc-104(R9Q)* mutant animals was only 58% of the WT adults, but the five suppressors restored the body length of *unc-104(R9Q)* to 72%-94% of the WT level. Notably, T102I and Y53F showed the highest rescue effects, while G265R displayed the least (Fig. 1D and F). The *unc-104(R9Q)* mutants exhibited severe coiling and curled postures, with limited spontaneous body elongation (Fig. 1D and G). We quantified the body curvature using a curling index (Fig. S1B and Methods), which was 0.36 ± 0.19 in *unc-104(R9Q)* mutant comparing to 0.08± 0.03 in WT, and found that all five suppressors significantly rescued the curling index, ranging from 0.08 ± 0.05 (Y53F) to 0.10 ± 0.06 (G265R) (Fig. 1D and G, and Fig. S1C). These results indicate that the intragenic mutations identified within the UNC-104 motor domain effectively rescue the loss-of-function caused by *unc-104(R9Q)* mutations, and their rescuing effects correlate with animal motility, body length, and morphology.

### Biochemical impacts of suppressors on the KIF1A nucleotide-binding pocket

In order to investigate the effects of the intragenic suppressors on the KAND R11Q mutation in vitro, we examined the motility of purified mouse KIF1A proteins as demonstrated previously^31^. The R11Q mutation is located within the nucleotide-binding pocket of the KIF1A motor domain, which is responsible for capturing ATP/ADP molecules (as depicted in Fig. 2A). In the WT motor domain, the arginine (R11) sidechain forms a hydrogen bond with the threonine (T106) sidechain (corresponding to T102I in UNC-104) at the bottom of the nucleotide-binding pocket. These residues wrap around the adenine ring of ATP/ADP, potentially facilitating nucleotide binding (Fig. 2A). Additionally, the tyrosine (Y56) residue (corresponding to Y53F in UNC-104) is located adjacent to R11, and its sidechain forms an additional hydrogen bond with the aspartic acid (D76) sidechain, which likely stabilizes the nucleotide-binding pocket (Fig. 2A).

**Fig.2.**
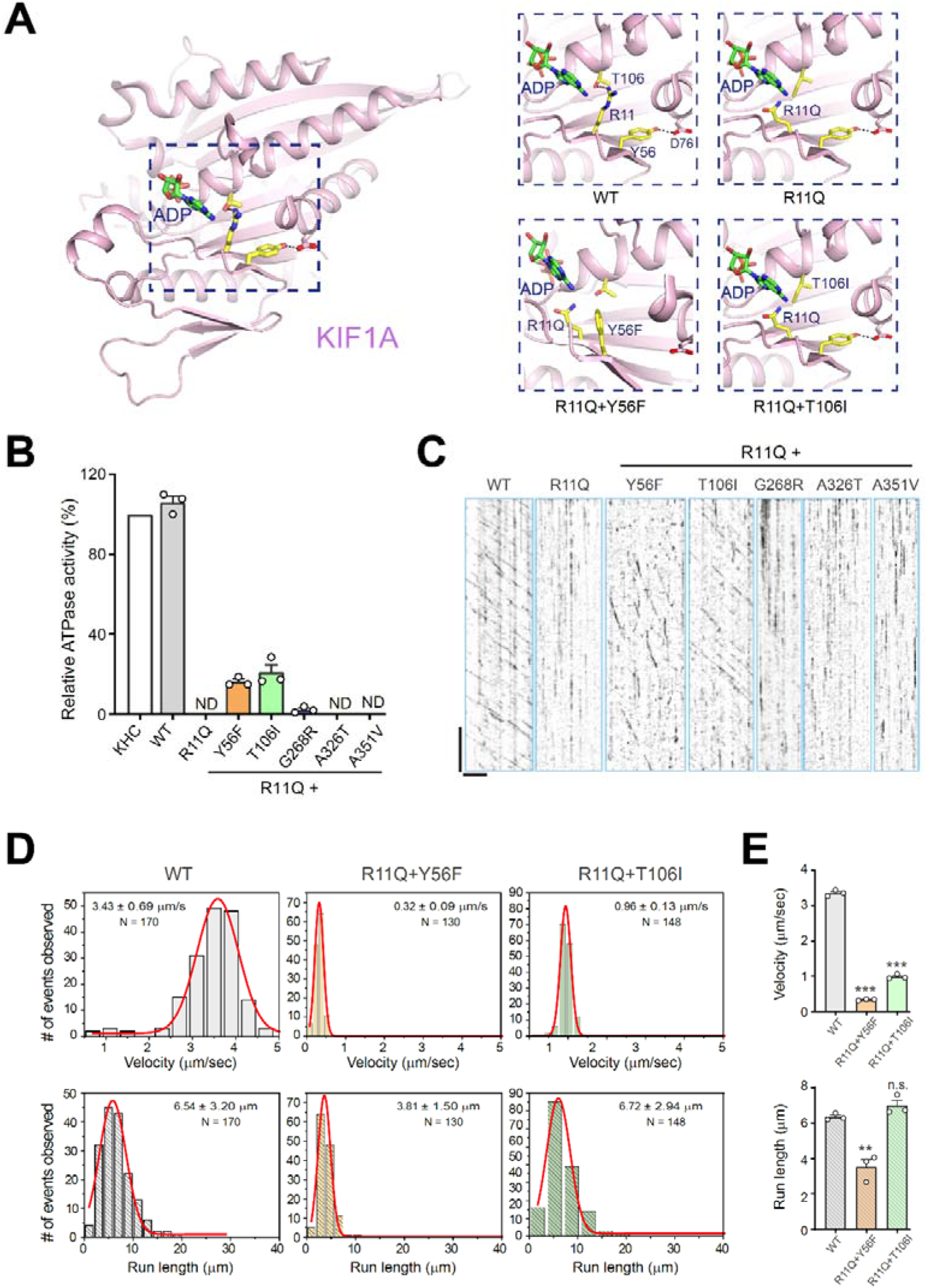
The structure, ATPase activity assay, and single-molecule study of KIF1A and mutants. (A) Potential impacts of mutations on the nucleotide-binding pocket in the motor domain of KIF1A (PDB code: 2ZFI). (B) Microtubule-stimulated ATPase activities of the wild-type and various mutants of KIF1A 1-613. Data were normalized by the microtubule-stimulated ATPase activity of kinesin-1 heavy chain (KHC) as 100%. Each experiment was repeated three times independently. Each bar represents the mean value ± SD, N.D., not detected. (C) Representative kymographs showing the motion of wild-type and various mutants of KIF1A 1-613 along the microtubule. Horizontal scale bar 5 µm; vertical bar 10 s. (D) Qualifications of the velocity (top) and run length (bottom) for each protein. The number of events is plotted for the velocity and run length as a histogram and fitted to a Gaussian distribution. The velocity and run length of the corresponding population of the protein (N) are indicated in each panel as mean ± SD. (E) Statistical analyses of the velocity (top) and run length (bottom) for each protein. Statistical significance for the velocity or run length is determined using Student’s t-test (n.s., not significant; ** *p* < 0.01; **** *p* < 0.0001) as compared with the WT, n=3 assays.

However, in the R11Q mutant, the substitution of the long-sidechain arginine with a short-sidechain glutamine may disrupt the hydrogen bonding interaction with T106 and hinder the packing with the adenine ring of ATP/ADP (Fig. 2A). Consequently, this mutation destabilizes the binding of nucleotides and deactivates the kinesin motor. Consistent with the structural analysis and previous studies ^13, 26^, the R11Q mutation abolished both the ATPase activity and processive movement of the motor domain of KIF1A (Fig. 2B-C).

Among the mutants that restored functionality, the Y56F and T106I substitutions in the nucleotide-binding pocket were able to restore the functionality of the R11Q mutation. In the R11Q and Y56F double mutant KIF1A, the removal of the hydroxyl group from Y56 disrupts the hydrogen-bonding interaction with D76. However, the bulky aromatic sidechain of the Y56F mutant, which is originally oriented towards the exterior of the pocket, tends to flip inside and occupy the void left by the R11Q mutation in the nucleotide-binding pocket of KIF1A (Fig. 2A). This flipping tendency likely compensates for the defect caused by the R11Q mutation. In the R11Q and T106I double mutant KIF1A, the substitution of the short-sidechain threonine with a long-sidechain isoleucine in the nucleotide-binding pocket also fills the vacancy created by the R11Q mutation. The T106I substitution forms additional contacts with the adenine ring of ATP/ADP, compensating for the loss of nucleotide binding (Fig. 2A). Therefore, both the Y56F and T106I substitutions in the R11Q mutant KIF1A structurally counteract the mutation-induced defect in the nucleotide-binding pocket.

In our biochemical assays, the Y56F and T106I mutations were found to partially restore both the ATPase activity and processive movement of the motor domain of KIF1A, as indicated by the velocity and processivity measurements (Fig. 2B-E). Notably, while the velocity remained compromised, the T106I mutation fully restored the run length of the motor domain (Fig. 2D-E), highlighting the positive impact of this substitution in renovating the nucleotide-binding pocket.

It is important to note that three other suppressor mutations, A319T, A344V, and G265R, which are not located in the vicinity of the nucleotide-binding pocket, did not exhibit any significant recovery of KIF1A(R11A) activity in vitro (Fig. 2D-E). This suggests that this group of mutations suppresses the animal UNC defects through mechanisms that are distinct from those of Y56F and T106I, which directly impact the nucleotide-binding pocket.

### Restoration of synaptic vesicle distribution and motility in suppressors

In our study, we examined the expression pattern and dynamics of UNC-104::GFP in *C. elegans*. In WT animals, we observed UNC-104::GFP fluorescence throughout the nervous system (Fig. 3A), which is consistent with previous findings and CeNGEN single-cell RNA-seq datasets^32^. However, in animals carrying the UNC-104(R9Q) mutation, we observed a significant reduction in GFP fluorescence intensity, particularly in the nerve ring region (Fig. 3A). Although the transcription (Fig. S2A) and expression pattern of UNC-104(R9Q) did not show a significant change, the overall fluorescence intensity was diminished.

**Fig.3.**
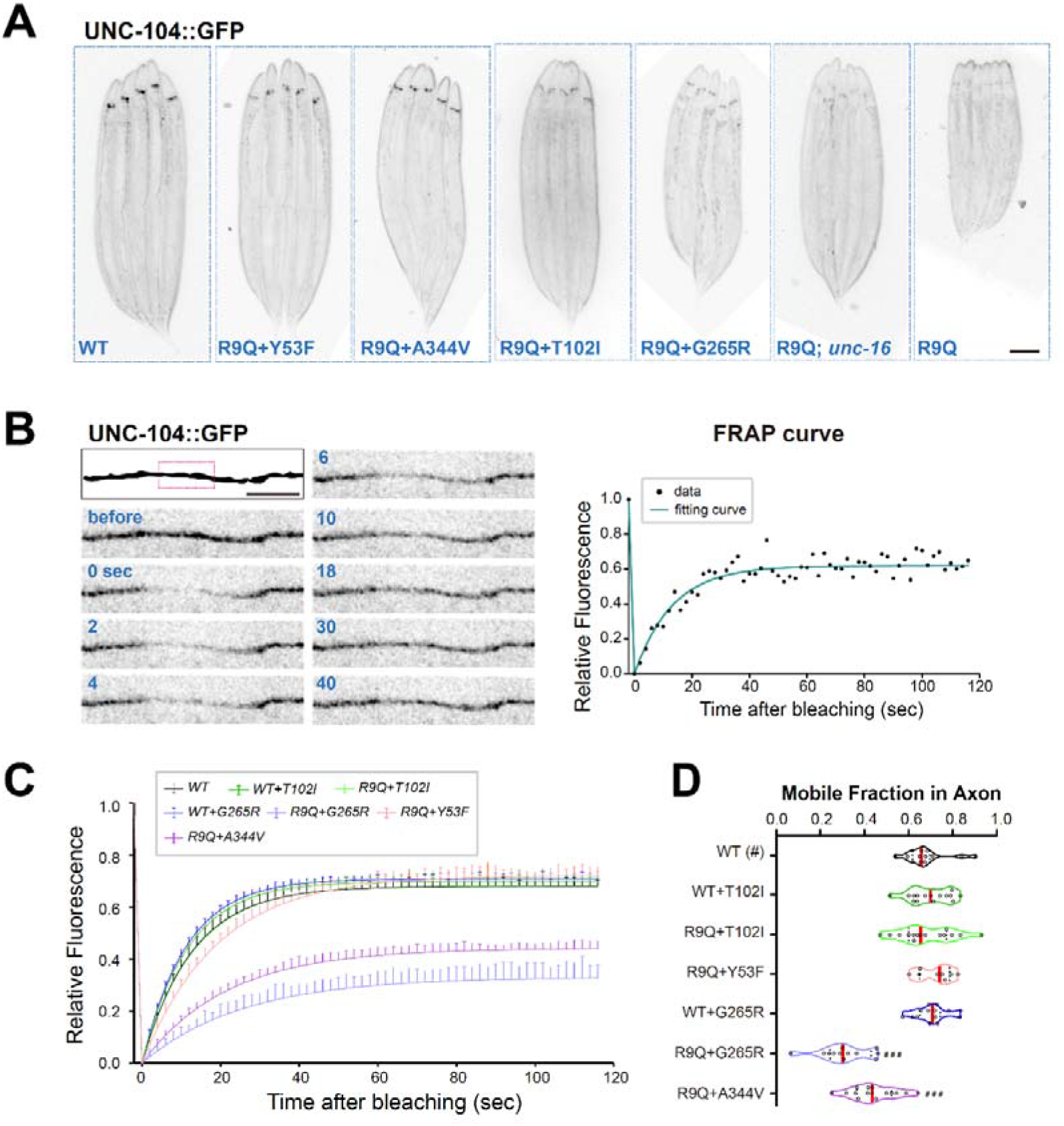
UNC-104::GFP dynamics in WT and mutants monitored by FRAP (A) Representative fluorescence images of mutation carrying UNC-104::GFP. GFP was knocked into the C-terminal of UNC-104 before R9Q mutation was introduced, and the *unc-104* mutations are indicated in each panel. Note that the fluorescence at the nerve ring was dramatically diminished in *unc-104(R9Q)* mutant, and this reduction was rescued to varying degrees in suppressors. Scale bar: 100 _μ_m. (B) UNC-104::GFP in axons was photobleached at the boxed area in the top left schematic panel, and recovery was recorded for two minutes. Images before and after photobleaching (at 0 second) at indicated time points are shown in sequential. Scale bar: 5 _μ_m. The kinetics of FRAP recovery were fitted with a single exponential equation. (C-D) Quantifications of *in vivo* UNC-104::GFP FRAP from different variants, showing that suppressors rescued the UNC-104(R9Q)::GFP kinetics (C) and the mobile portion (D) in the axon. Error bars represent SEM, n = 8 to 15 independent animals. Statistical significance was calculated by the unpaired Student’s *t*-test: ^#^ ^#^ ^#^ *p* < 0.001 compared to the WT animals. See also Fig. S1.

Remarkably, the intragenic suppressors were able to substantially restore the GFP intensity, indicating their rescuing effects on UNC-104(R9Q) function (Fig. 3A). To further investigate the dynamics of UNC-104, we conducted fluorescence recovery after photobleaching (FRAP) experiments in the dorsal nerve cords (Fig. 3B, outlined in red). In UNC-104(R9Q) animals, the GFP fluorescence was barely detectable in this region, whereas the genetic suppressors restored GFP fluorescence (Fig. S2B-C).

Two of the suppressors, UNC-104(R9QY53F) and UNC-104(R9QT102I), showed similar GFP mobile ratios, indicating the recovery of dynamic behavior comparable to the WT motor (Fig. 3B-D). The GFP recovery ratios for these suppressors were 0.72 ± 0.08 and 0.70 ± 0.10, respectively, which were in the range of the WT motor (0.68 ± 0.10, mean ± SD) (Fig. 3B-D and Fig. S2B). On the other hand, UNC-104(R9QA344V) and UNC-104(R9QG265R), which exhibited partial rescue effects on animal movement (Fig. 1E), also partially restored the GFP recovery ratio (Fig. 3C-D and Fig. S2C). These findings provide further evidence of the rescuing effects of intragenic suppressors on the cellular dynamics of UNC-104 and support their correlation with the partial restoration of animal movement observed in our study.

To investigate the distribution of synapses in different genetic backgrounds, we focused on the DA9 cholinergic motor neuron in the *C. elegans* dorsal nerve cord (DNC). In WT animals, the DA9 neuron typically forms approximately 25 en-passant synapses along a proximal axon segment in the DNC (Fig. 4A)^29^. We visualized these synapses using a transgenic strain expressing the synaptic vesicle protein (SVP) marker mScarlet::RAB-3^7, 26, 33^ under the control of the DA9 promoter (P*mig-13*). In *unc-104(R9Q)* mutant animals, the DA9 neurons showed a severe deficiency in en-passant synapses (Fig. 4A-C). The length of the synaptic region in DA9 neurons was significantly reduced from 115.10 ± 12.69 μm in WT animals to 20.48 ± 26.09 μm in *unc-104(R9Q)* animals (Fig. 4A-B). Furthermore, the number of mScarlet::RAB-3 puncta, which represents individual synapses, dramatically decreased from an average of 28.36 ± 5.77 in WT animals to only 2.08 ± 2.04 in *unc-104(R9Q)* animals (Fig. 4A and C). Additionally, the size and intensity of mScarlet::RAB-3 puncta were also reduced in *unc-104(R9Q)* animals (Fig. 4D-E). Importantly, the intragenic suppressors were able to significantly restore the distribution area, number, and size of synapses in the DA9 neuron (Fig. 4A-E). These findings suggest that the identified suppressor mutations have a rescuing effect on the synaptic defects caused by the *unc-104(R9Q)* mutation.

**Fig.4.**
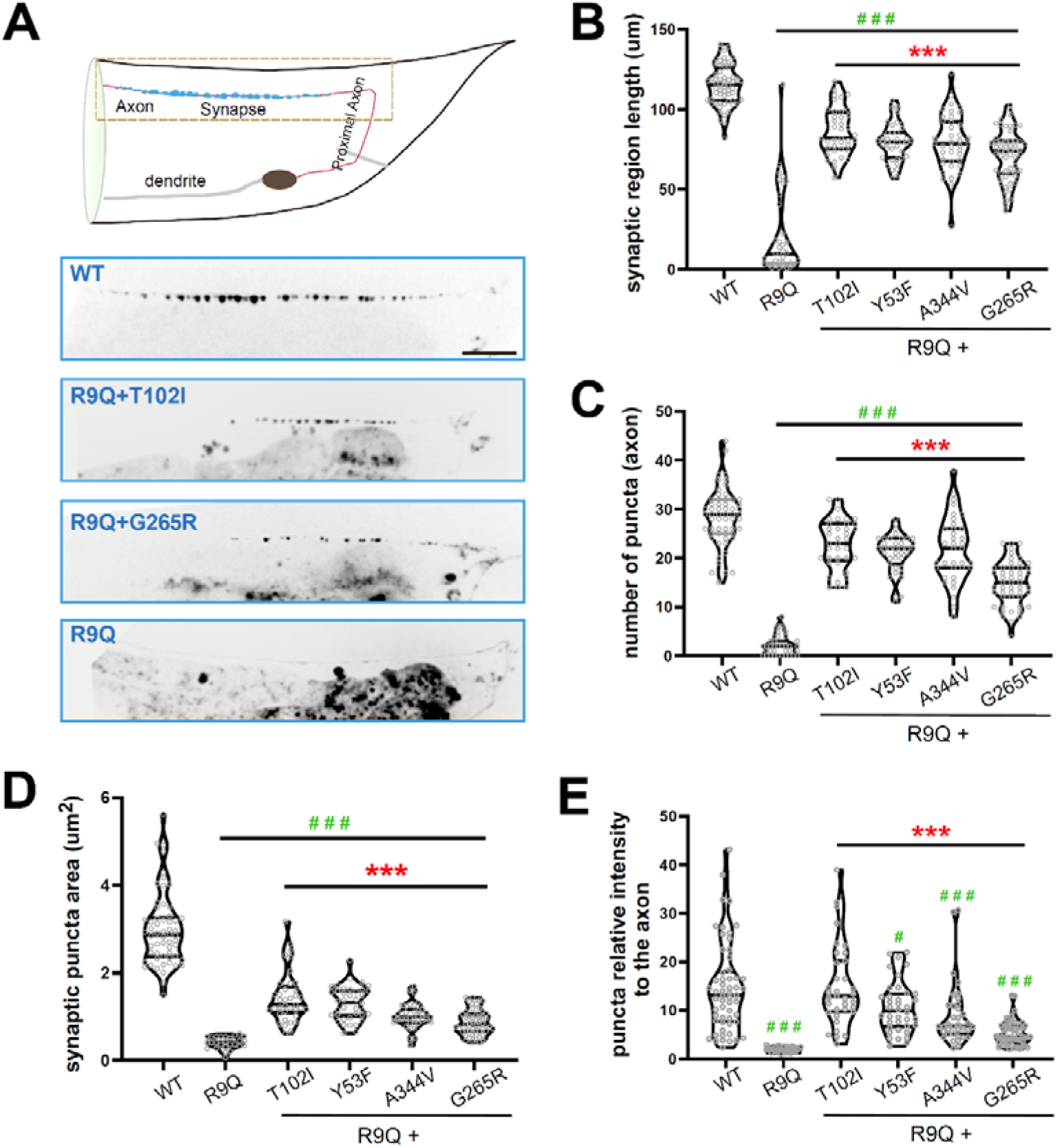
Distribution and morphology of synaptical vesicles in WT and mutant *unc-104* animals. (A) Schematic diagram of the DA9 neuron. The dorsal axon region framed with a dashed line was imaged in each genotype as shown in the panels below for synaptic vesicle analysis. Blue dots along the axon represent synaptic vesicles. The proximal axon was used for velocity analysis in Fig.5. Representative images show the distribution of synaptic vesicles, visualized by P*mig-13*::mScarlet::RAB-3, in the DA9 neuron in WT, *unc-104(R9Q)* mutant and rescuers. Scale bar: 10 _μ_m. (B-E) Violin plots with raw data dots showing the quantifications of the synaptic puncta characters including the length of synaptic region (B), number of puncta (C), puncta size (D), and relative fluorescence intensity normalized to axon regions without synaptic vesicles (E), in the dorsal axon of WT, *unc-104(R9Q)* mutant and suppressors. n = 39 to 60 independent animals. Statistical significance was calculated by the unpaired Student’s *t*-test: *** *p* < 0.001, compared to the *unc-104(R9Q)* mutant; ^#^ *p* < 0.5, ^#^ ^#^ *p* < 0.01, ^#^ ^#^ ^#^ *p* < 0.001 compared to the WT animals.

To investigate whether the axonal synaptic defects in unc-104(R9Q) mutants were attributed to vesicle transport impairments, we examined the motility of mScarlet::RAB-3 puncta in the proximal region of the DA9 axon (Fig. 4A)^33^. By performing kymography analyses, we were able to track and analyze the movement of RAB-3 puncta in both anterograde and retrograde directions. In WT animals, we observed frequent axonal transport of RAB-3 puncta in both anterograde and retrograde directions (Fig. 5A and E-F). However, in *unc-104(R9Q)* mutants, anterograde RAB-3 transport was disrupted, and there was a significant reduction in retrograde transport events (Fig. 5A and E-F), indicating impaired motility of UNC-104 along the axon. Remarkably, the intragenic suppressors were able to restore the frequency of anterograde and retrograde transport events, as well as the distribution and motility of synaptic vesicles in *unc-104(R9Q)* mutant animals (Fig. 5A-G). This restoration of vesicle transport dynamics explains the recovery of animal UNC phenotypes observed in these suppressors. Notably, the A344V and G265R suppressors, which rescued animal movement but did not restore the motor activity of UNC-104(R9Q) *in vitro* (Fig. 2B-C), partially recovered the synaptic defects associated with UNC-104’s function in *C. elegans* (Fig. 5A-G, and Fig. S3). This suggests that A344V and G265R likely employ distinct rescuing mechanisms from those of Y56F or T106I. Overall, our findings indicate that the intragenic suppressors not only rescue animal motility and morphology, but also restore the synaptic defects and dynamics of synapses in the DA9 cholinergic motor neuron. These results further support the functional significance of the identified suppressor mutations and suggest that targeting some of these sites within the UNC-104/KIF1A motor domain may have potential therapeutic implications for KAND.

**Fig.5.**
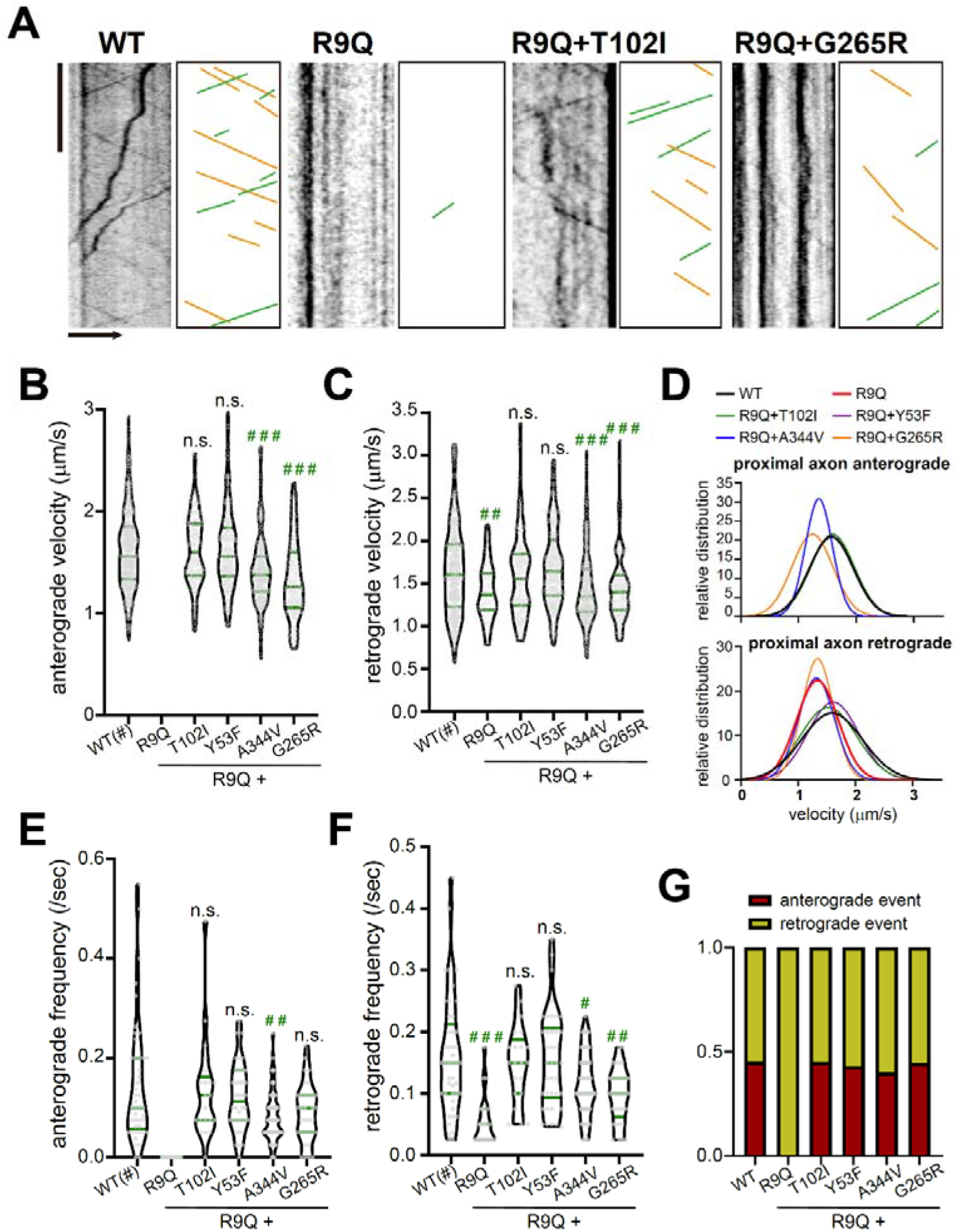
Motility of synaptical vesicles in WT and mutant *unc-104* animals. (A) Representative kymographs depicting the axonal anterograde (left-to-right) and retrograde movement of synaptic vesicles labeled with mScarlet-tagged RAB-3 in WT, *unc-104(R9Q)*, *unc-104(R9QT102I* and *unc-104(R9QG265R)* worms. Processive transport events in the original kymograph are marked with lines in the schematic diagram. Scale bars: horizontal represents 5 _μ_m, arrow pointing towards distal axon, and vertical represents 10 s. (B-C) The velocity of axonal anterograde transport (B) and retrograde transport (C) are plotted as violin graphs with raw data dots, where the green bars indidate the median and quartiles. At least 10 independent worms of each genotype were analyzed. (D) Overlay of Gaussian distribution fitting plots of anterograde (top) and retrograde (bottom) synaptic vesicles transport velocities. See also Fig. S2. (E-F) The frequency of axonal anterograde transport (E) and retrograde transport (F) are plotted as violin graphs harboring raw data dots, with the green bars indicating the median and quartiles. At least 10 independent worms of each genotype were analyzed. (G) The ratio between the directionality of vesicle movement is shown in the bar graph. Statistical significance was calculated by the unpaired Student’s *t*-test: n.s. not significant, ^#^ *p* < 0.5, ^#^ ^#^ *p* < 0.01, ^#^ ^#^ ^#^ *p* < 0.001 compared to the WT animals.

### Fisetin alleviates animal movement and morphological defects in UNC-104(R9Q)

Having established the capacity to rescue UNC-104(R9Q) abnormalities through several distinct pathways *in vivo*, we aimed to explore the potential of small-molecule chemicals in restoring animal behavior. Our accidental starvation of *C. elegans* carrying the UNC-104(R9Q) mutation led to intriguing observations of improved body morphology and animal movements (Fig. S4A-B), which suggests that nutrient conditions may play a role in modulating the behaviors and phenotypes associated with the mutation. This inspired us to explore the potential effects of commercialized food supplements known for their benefits to the nervous system on UNC-104(R9Q) animals. Our initial focus was on fisetin^34, 35^ (Fig. S4C), a plant flavonol, due to its availability, low cost, and neuroprotective properties. Fisetin has shown effectiveness in improving behavior and physiology in aging mice, preserving cognitive function in Alzheimer’s disease models, and promoting survival in *C. elegans* exposed to thermal stress ^36–39^. We found that supplementing fisetin in the growth media of *C. elegans* resulted in improved motility and alleviation of developmental and behavioral defects in *unc-104(R9Q)* mutant animals (Fig. 6A). The effects were dose-dependent, with higher doses showing greater improvements in motility, body length, and posture (Fig. 6A-D). Additionally, we observed a partial rescue of both the distribution and quantity of neuronal synapses following fisetin treatment (Fig. 6E-F). This implies that the amelioration of animal behavioral abnormalities may be attributed to the restoration of synaptic functionality. These results underscore the potential of fisetin supplementation as a promising therapeutic strategy for alleviating the symptoms associated with KAND.

**Fig.6.**
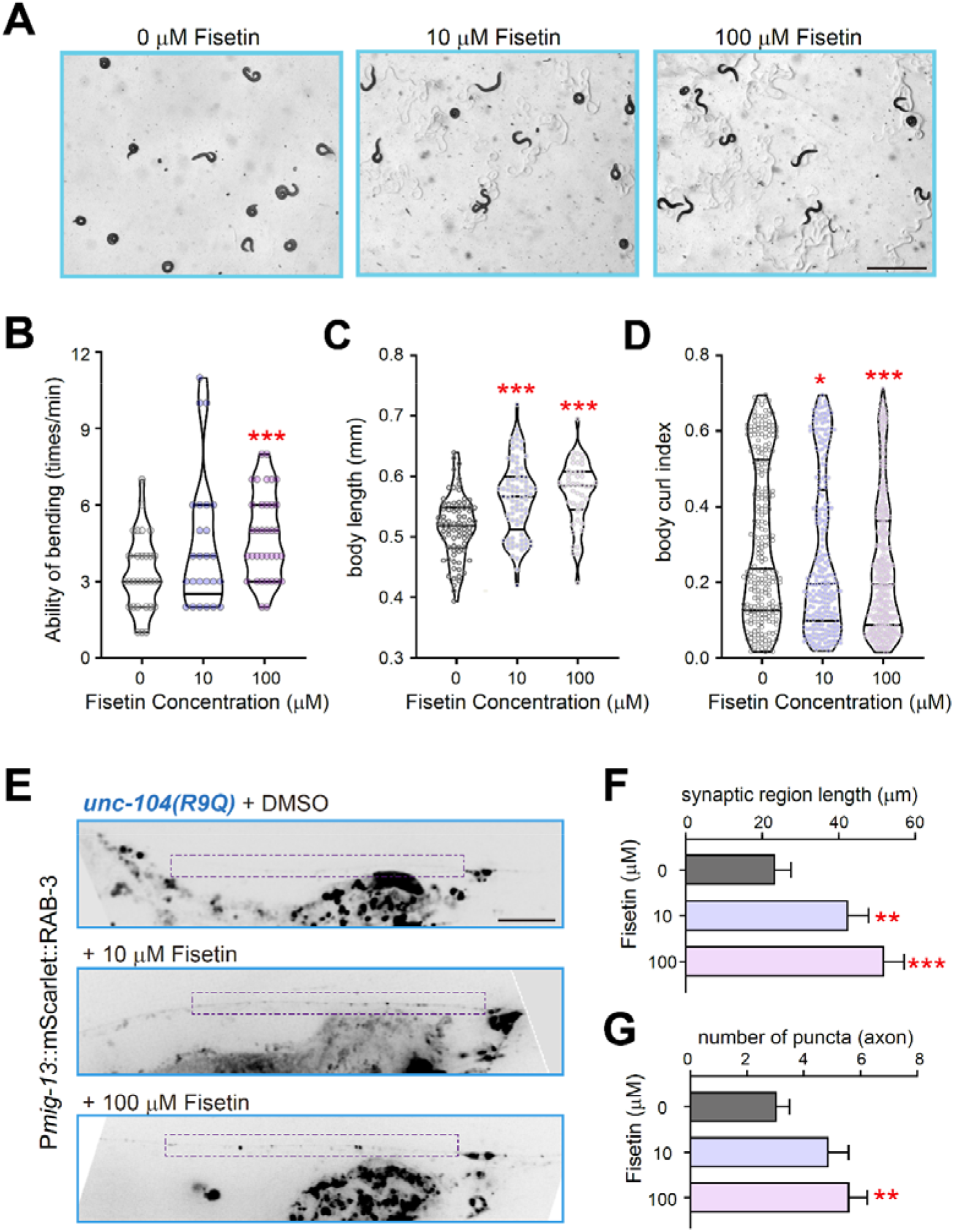
Fisetin partially recovered the motility and body morphology of *unc-104(R9Q)* animals. (A) Representative bright field images of *unc-104(R9Q)* animals treated with different concentrations of Fisetin. (B-D) Violin plots harboring raw data dots showing the quantification of animal swimming behavior (B, n = 25-30 animals), body length (C, n = 45-85 animals), and curl index (D, n =230-360 animals) morphology phenotypes of *unc-104(R9Q)* animals treated with different concentrations of Fisetin. Green bars represent the median and quartiles. Statistical significance was calculated by the unpaired Student’s *t*-test: * *p* < 0.05, *** *p* < 0.001, compared to the *unc-104(R9Q)* animals cultured on the control NGM plates. (E) Representative images show the distribution of synaptic vesicles, visualized by P*mig-13*::mScarlet::RAB-3, in the DA9 neuron in *unc-104(R9Q)* mutant with indicated drug treatments. Scale bar: 10 _μ_m. (F-G) Bar graphs showing the quantifications of the length of synaptic region (F) and number of puncta (G) in the dorsal axon of *unc-104(R9Q)* mutant with indicated drug treatments. n = 40-50 independent animals. Statistical significance was calculated by the unpaired Student’s *t*-test: ** *p* < 0.01, *** *p* < 0.001.

### Fisetin directly restored the motor activity of KIF1A(R11Q) *in vitro*

The observed mitigating effects of fisetin on the UNC-104(R9Q) animal prompted us to investigate its potential direct impact on the KIF1A protein. To assess this, we first perform docking analysis between fisetin and the KIF1A motor with the R11Q mutation (Fig. 7A). The docking results revealed the effective binding of fisetin to the adjacent pocket of Q11 in the mutant structure, likely restoring the structural stability of the region surrounding Q11, which was compromised by the R11Q mutation. These findings provide valuable structural insights into the observed functional rescue of the R11Q mutant upon fisetin treatment. Conversely, in the WT KIF1A motor domain, the presence of the R11 guanidine group partially occupied the pocket, thereby hindering the binding of fisetin (Fig. 7A, right panel). These results underscore the specific recognition and functional rescue of KIF1A-R11Q by fisetin without affecting the wild-type KIF1A motor.

**Fig.7.**
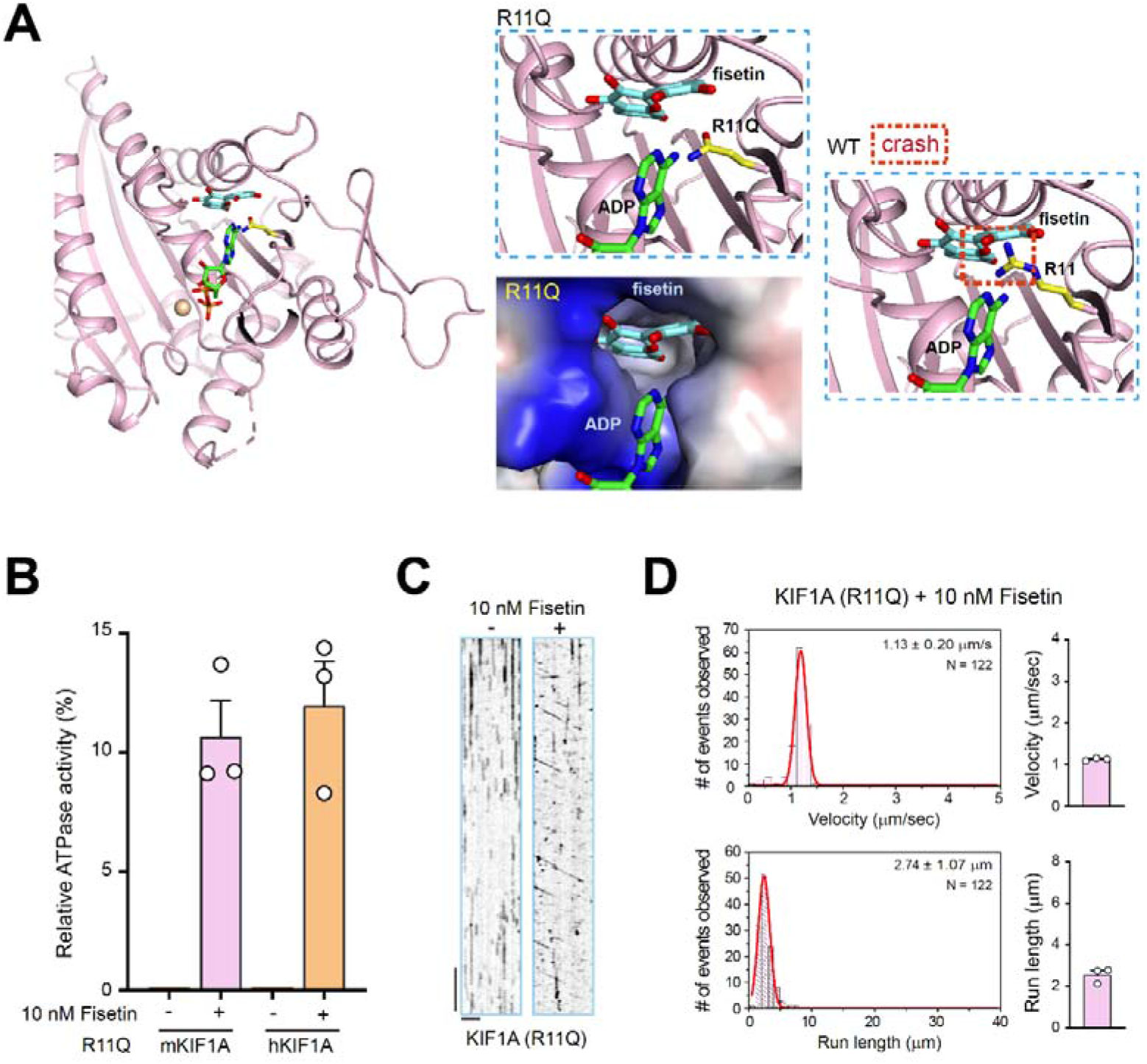
Fisetin restored the motor activity of KIF1A(R11Q) *in vitro*. (A) Structural model of fisetin docked into the pocket in the KIF1A(R11Q) mutant. Right panel, guanidine group of Arg11 clashes with fisetin. KIF1A PDB code: 2ZFI. (B) Microtubule-stimulated ATPase activities of the human and mouse KIF1A 1-613 carrying R11Q mutation, with or without 10nM fisetin. Data were normalized by the microtubule-stimulated ATPase activity of kinesin-1 heavy chain (KHC) as 100%. Each experiment was repeated three times independently. Each bar represents the mean value ± SD. (C) Representative kymographs showing the motion of mouse KIF1A(R11Q) 1-613, with or without 10nM fisetin, along the microtubule, respectively. Horizontal scale bar 2 µm; vertical bar 5 s. (D) Qualifications of the velocity (top) and run length (bottom) of mouse KIF1A(R11Q) protein with 10nM fisetin treatment. The number of events is plotted for the velocity and run length as a histogram and fitted to a Gaussian distribution, and the average of three repeats were plot as bar graph on the right. The velocity and run length of the corresponding population of the protein (N) are indicated in each panel as mean ± SD.

Subsequently, we generated recombinant mouse and human KIF1A(R11Q) proteins (Fig. S5A) and conducted biochemical analyses to assess the impact of fisetin on its ATPase activity. We observed negligible ATPase activity in KIF1A(R11Q) treated with 0.1% DMSO, the solvent used for fisetin. However, the presence of 10 nmol/L fisetin led to the detection of ATPase activity (Fig. 7B). Although fisetin only increased the ATPase activity of KIF1A(R11Q) to approximately 12% of that in WT KIF1A, the intragenic suppressors Y57F and T102I, which effectively rescued the UNC phenotype caused by R11Q, elevated the ATPase activity of KIF1A(R11Q) to 15% and 22% of the WT level, respectively (Fig. 2B). This suggests that even a modest restoration of ATPase activity is potentially sufficient to rescue behavioral defects at the animal level.

Furthermore, we performed a microtubule-gliding assay to examine whether fisetin treatment could induce force generation in KIF1A(R11Q). In the absence of fisetin, KIF1A(R11Q) did not exhibit any MT gliding in vitro. However, the addition of 10 nmol/L fisetin to the gliding assay enabled MT to glide at a speed of 0.49 μm/s, which is comparable to that of WT KIF1A (Fig. S5B and Movie S3). Additionally, using a single-molecule assay, we demonstrated that fisetin treatment enabled KIF1A(R11Q) to generate processive movement along MT, although the speed and run length were lower than those of WT KIF1A (Fig. 7C-D, and Movie S1-2). As predicted by molecular docking, we did not observe any effects of fisetin on the ATPase activity or processive movement of the wild-type KIF1A (Fig. S5C). These findings indicate that fisetin specifically enhances the functionality of the KIF1A(R11Q) mutant protein while exerting no effects on the wild-type KIF1A.

## Discussion

Our unbiased genetic suppressor screens have successfully identified 20 mutations located within the motor domain of UNC-104 that can rescue the detrimental effects caused by the KAND-associated amino acid substitution. Through comprehensive biochemical and single-molecule studies, we have classified these suppressor mutations into two distinct groups. The first group, represented by Y56F and T106I, is situated in the vicinity of the nucleotide-binding pocket. These mutations restore the ATP/ADP binding capacity of UNC-104(R11Q), leading to the recovery of its ATPase activity and motility. In contrast, the second group of suppressors, including A319T, A344V, and G265R, does not exert any influence on the nucleotide-binding pocket and fails to restore the biochemical and biophysical properties of UNC-104(R11Q) *in vitro*. Despite the divergent effects observed *in vitro*, both sets of amino acid substitutions have demonstrated their efficacy in rescuing the abnormalities in synapse distribution and animal behavior associated with the R11Q mutation. This underscores the remarkable potential for restoring function even in the presence of a deleterious KAND mutation that completely abolishes the motor’s ATPase activity. Furthermore, the involvement of multiple distinct mechanisms in the restoration process provides great inspiration and suggests that effective interventions for KAND are not only feasible but can be achieved through diverse strategies.

The suppressors belonging to group one suggest that if a secondary mutation can counterbalance the structural defects induced by the primary mutation, it has the potential to restore motor activity. In other words, it may be possible to rescue the defects resulting from the primary mutation even without directly correcting it, as long as a second mutation can counteract its structural effects. This concept carries significant implications for drug development. A small molecule that mimics the effects of the Y56F or T106I substitution in the nucleotide-binding pocket could potentially restore the motor activity of UNC-104(R11Q). Importantly, the T106I single mutation does not result in any detectable abnormalities in *C. elegans*, as shown in Figure 1E. Therefore, we propose that such a small molecule-based intervention strategy may not impact the normal function of WT UNC-104/KIF1A but specifically improve the R11Q mutant.

While we have identified several independent mutant alleles of Y56F or T106I, we suspect that our suppressor screens are not yet saturated due to the limitations of the chemical mutagen ethyl methanesulfonate (EMS) used in this study. EMS predominantly induces G/C to A/T nucleotide conversions, thereby only targeting a limited subset of codons^40–42^. Future investigations will explore alternative chemical mutagens, such as ENU, to introduce a broader spectrum of mutations for selection. Hence, we speculate that Y56F and T106I are not the sole targets for drug development. As long as the nucleotide-binding pocket can be restored through the substitution of other residues or by utilizing small molecules, the defects caused by the R11Q mutation can be remedied. Given that mutations in the motor domain of the kinesin family can contribute to various genetic disorders, including those beyond KAND, our design principles hold implications for addressing other kinesin-associated pathologies^1, 3, 43, 44^.

Equally intriguing are the suppressors in the second group, which effectively restore synaptic and behavioral defects in vivo but do not exhibit restored activity of R11Q in vitro. Considering our current knowledge of motor regulation, it is challenging to conceive a model that can reconcile the differences observed between *in vitro* and *in vivo* outcomes. While we have not yet validated all the substitutions in this group as bona fide suppressors, we conducted double knock-in experiments that generated G265R and R11Q double mutant animals *de novo*, thereby confirming the suppressive effect of G265R on R11Q. The acquisition of multiple alleles through independent screens for many suppressors suggests that we may have identified causative agents capable of restoring R11Q function. These suppressors are likely to act through diverse mechanisms, such as influencing the conformational stability of the protein, modulating the interaction between the motor domain and its cargo, or impacting the recruitment of the motor to the microtubule track. We also propose the existence of previously unknown factors harboring suppressive properties within the second group, which could play a crucial role in activating UNC-104(R11Q), considering the involvement of various intracellular factors that either activate or inhibit motor activity^44–47^. One plausible scenario is that an intracellular factor binds to a motif containing G265 and inhibits wild-type UNC-104, and the G-to-R substitution disrupts this interaction, thereby releasing the inhibition *in vivo*. Alternatively, the G-to-R substitution may lead to the ectopic recruitment of an axonal protein to interact with UNC-104(R11Q), thereby enhancing its motor activity within axons. Future genetic screens involving UNC animals in the background of these rescued animals could potentially shed light on the identification of such molecular candidates. Nonetheless, we assert that motor regulation is likely to be more intricate and fascinating than previously believed, and the exploration of clinical motor mutations will unveil new avenues for motor research.

Our study emphasizes the significance of unbiased genetic screens in unraveling the complexities of diseases and devising effective interventions. Although artificial intelligence (AI) has transformed biomedical research by generating numerous hypothetical models, it is noteworthy that AI cannot predict the genetic suppressors identified in this study, despite the availability of KIF1A’s structural information. Artificial evolution, on the other hand, which relies on extensive unbiased genetic screens and whole-genome sequencing, holds great promise in complementing AI and providing novel insights into biology and the understanding of diseases.

The unexpected and inspiring direct effects of fisetin on the UNC-104(R11Q) protein shed new light on its therapeutic potential. The kinesin family motor protein has been a subject of drug development efforts over the past decades; however, the efficacy of developed kinesin antagonists or inhibitors has posed limitations for therapeutic interventions, despite their widespread use in laboratories^48^. Our findings on fisetin suggest that the development of agonists targeting disease-associated kinesins harboring human genetic mutations could be a fruitful and valuable future direction for motor protein-based drug discovery.

We acknowledge that our study on fisetin in UNC-104(R11Q) animals is still in its preliminary stages. Further experimentation is required to determine the exact interaction between KIF1A and fisetin, and the observed effects of fisetin on improving animal movement or morphology are modest compared to our genetic suppressors. Additionally, fisetin may regulate various complex processes that contribute to the overall physiological improvement in animals. Therefore, we propose considering fisetin as a food supplement rather than a drug for a specific subgroup of KAND patients. Due to its chemical properties and diverse biological effects, fisetin alone may not be an ideal drug; however, the use of this small molecule, along with the proof-of-concept and design principles demonstrated in our study, can provide valuable insights for the development of specific and potent drugs targeting KAND. Considering the reproducible effects of fisetin in both in vivo and in vitro settings, as well as its cost-effectiveness, the application of fisetin as a nutrient supplement may offer potential benefits to KIF1A(R9Q)-associated KAND patients. Furthermore, given the potential renovating effects of fisetin on the nucleotide binding pocket, it cannot be excluded that fisetin may also improve KIF1A carrying other mutations that affect this region.

Caution is crucial when considering the use of fisetin or any other dietary supplement for KAND patients. Safety considerations, including appropriate dosage, potential interactions with other medications or underlying conditions, should be carefully assessed. We strongly recommend consulting with healthcare professionals or experts in the field of neurodevelopmental disorders before initiating any fisetin supplementation regimen for individuals with KAND.

## Materials and Methods

### Strains and Genetics

*C. elegans* strains were maintained as described previously^49^, on nematode growth medium (NGM) plates with OP50 feeder bacteria at 20 °C. To administer fisetin (Sigma, Cat# 528-48-3), the drug was added to the NGM medium at the desired concentration prior to pouring the plates, which were subsequently seeded with OP50 culture in the usual manner. All the engineered *C. elegans* strains were genetic derivatives of the strain Bristol N2, Strains used in this study is summarized in ***supplementary table S2***. The PHX4787 (UNC-104::GFP) strain, referred as wild-type in this study, and other CRISPR/Cas9-mediated genome editing strains were created by Suny Biotech (Fuzhou City, China). The detailed information such as modified locus is listed in ***supplementary table S3***. Transformation of *C. elegans* to introduce the P*mig-13::mScalet::RAB-3* was performed by DNA injection as described^50^, and the information of plasmids and primers are described in ***supplementary table S4.*** All animal experiments were performed following governmental and institutional guidelines.

### Suppressor screening

EMS mutagenesis was performed as described before^51, 52^ with some modifications. PHX5261 [*unc-104(R9Q)::GFP*] worms were cultured on regular NGM plates seeded with OP50 feeder till the population was just starved. Then the mixed-stage worms from 10-20 plates were collected, and L1-like larva were sorted with 15 μm diameter metal filter. The sorted larvae were cultured to late L4, harvested and treated with ethyl methanesulfonate (EMS). After the treatment, worms were washed and dispersed to 200-400 OP50-seeded NGM plates by dropping 5-10 worms each plate close to the plate margin. Plates were examined when the majority of F2 progenies were older than L4, and basically worms that were able to move to the opposite margin of the plates were very likely to be suppressors. Approximately 100,000 haploid genomes were checked after multiple runs of screens, and suppressors from different plates were considered independent. After phenotype confirmation over two generations, suppressors were sent to whole genome sequencing followed by comprehensive bioinformation analyses and genetic manipulations to identify and validate the genomic mutations responsible for the rescue.

### RNA-sequencing analysis

Adult worms fed on regular NGM plates were bleached to obtain mixed stage embryos, and the hatched larval were synchronized on NGM plates without OP50 for 16 hours before harvest. Total RNA was isolated with TRIzol reagent for RNA-Seq library construction and sequenced using on Illumina HiSeq-PE150 instruments. Two biological replicates were prepared for each library type. Raw sequencing reads were trimmed using TrimGalore, paired-end reads with at least 20 nucleotides in length were aligned to *C. elegans* reference genome cel235 using STAR and quantified by HTSeq. The RPKM of each gene was calculated based on the uniquely mapped gene read counts mapped to this gene.

### Imaging and analysis of the worm phenotypes

A scientific cMOS camera (IMX236, Sony Semiconductor Solutions Corporation) was mounted to a compound microscope for bright field images. Swim test was performed as described previously^25, 26, 30^ with day 1 young adult worms. Synchronized late L4 stage animals were imaged on NGM plates for body length and curvature analysis. The imaging processing, measurement and index calculation were described in Fig. S1B with ImageJ (http://rsbweb.nih.gov/ij/), and tightly curled animals were not taken into the body length analysis.

### Live-cell imaging

Synchronized late L4 *C. elegans* animals were mounted on 2% (wt/vol) agarose pads and anesthetized with 0.1 mmol/L levamisole in M9 buffer for live-cell imaging. Our imaging system includes an IX83 inverted microscope (Olympus Lifescience, Inc.) equipped with an EM CCD camera (Andor iXon+ DU-897D-C00-#BV-500), and the 488-nm and 561-nm lines of a Sapphire CW CDRH USB Laser System to a spinning disk confocal scan head (Yokogawa CSU-X1 Spinning Disk Unit). The UNC-104::GFP signal of whole animal were imaged with a 10X/0.3NA UPlan Semi Apochromat WD objective. The steady state images of mScarlet::RAB-3 fluorescent puncta along DA9 neuron axons were taken with 60X, 1.30NA UPlanSApo silicone oil immersion objective at an exposure time of 200 milliseconds, and time-lapse images showing mScarlet::RAB-3 transports were acquired with 100X, 1.49NA Apochromat oil immersion objective at an exposure time of 200 milliseconds without interval time for 200 frames. Kymograph extraction, image processing, and measurement of axonal synapse morphology and movement speeds were performed in ImageJ.

### FRAP

The fluorescence recovery after photobleaching (FRAP) experiments in *C. elegans* were carried out on the L4 larva worms with GFP reporter knocked into the C-terminal of UNC-104, using a laser scanning confocal microscope (LSM900, Zeiss) equipped with a 63x/1.4 NA Plan-Apochromat oil immersion objective. Photobleaching, image acquisition, and raw fluorescence measurement were performed on ZEN system under identical settings for all experiments. When conducting the assay, the puncta-free GFP-positive dorsal nerve cord processes were photobleached over a rectangular region (65LJ×LJ30LJpixels, 0.085 μm/pixel) using 40 iterations of the 488-nm laser with 10% laser power transmission and the pinhole diameter 200μm. Images were collected with 1% 488-nm laser power over 250LJ×LJ60LJpixels at 8-bit intensity resolution using a pixel dwell time of 0.96 μs. Prior to the bleaching, 2 frames of images were acquired, and the recovery of fluorescence was recorded for 120 s with 2 s interval to guarantee no further recovery was detected. After background subtraction, fluorescence recoveries were calculated as previously described^53^.

### Protein expression and purification

DNA sequences encoding mouse KIF1A fragment including the MD-NC-CC1-FHA tandem (residues 1–613), and various mutants were each cloned into a modified version of the pET32a vector. The human KIF1A protein is derived by mutating the corresponding residues in the mouse KIF1A protein to match their human counterpart, based on the amino acid sequence alignment (Fig. S5A). All the mutations in the KIF1A fragment were created using the standard PCR-based mutagenesis method and confirmed by DNA sequencing. Recombinant proteins were expressed in *Escherichia coli* BL21 (codon plus) host cells at 16 °C. After the 16 h cultivation, the cells were harvested and suspended in the buffer containing 50 mM Tris-HCl, pH 7.5, 500mM (NH4)_2_SO_4,_ 2mM MgCl_2_, 1mM EGTA, 1 mM PMSF. The proteins were purified by Ni^2+^-Sepharose 6 Fast Flow (GE healthcare) affinity chromatography with the washing buffer (50mM Tris-HCl, pH 7.5, 500mM (NH4)_2_SO_4_, 2mM MgCl_2_, 1mM EGTA, 40mM imidazole) and the elution buffer (50mM Tris-HCl, pH 7.5, 500mM (NH4)_2_SO_4_, 2mM MgCl_2_, 1mM EGTA, 500mM imidazole). The proteins were further purified by size-exclusion chromatography (Superdex-200 26/60, GE healthcare) with the buffer containing 150mM NaCl, 50mM Tris-HCl, pH 7.5, 2mM MgCl_2_, 1mM EGTA, 1mM DTT.

### Microtubule-stimulated ATPase assay

Measurements of the microtubule-stimulated ATPase activities of KIF1A MD-NC-CC1-FHA and various mutants were performed by using the HTS Kinesin ATPase Endpoint Assay Biochem Kit (Cytoskeleton, Inc., BK053). Briefly, all of the measurements were based on the malachite green phosphate assay to probe inorganic phosphate generated during the reaction. A standard curve of phosphate was made to estimate the amount of phosphate generated. Each protein sample had three replicates and each measurement was repeated at least three times independently. The kinesin-1 heavy chain (KHC) supplied in the Kit was used as the control. All of the data were analyzed by using the GraphPad prism8 program.

### Tubulin purification and labeling

Crude tubulins were obtained from porcine brain through double cycles of polymerization and depolymerisation, as previously described^54^. Tubulins were further purified using a TOG-based affinity column^55^. Tubulins were labeled with TAMRA (Thermo Fisher Scientific) using NHS esters, following standard protocols^56^.

### Polymerization of taxol-stabilized microtubules

A previously published protocol was followed for the *in vitro* motility assay^57^. Briefly, short microtubule seeds were prepared by incubating 32 μM porcine brain tubulin mixture containing 5% TAMRA-labeled tubulin with 1 mM GTP, 4 mM MgCl_2_ and 4% DMSO. After incubation on ice for 5 min, the mixture was polymerized in a 37 °C water bath overnight in the dark. Then, 400 μl of warm BRB80 buffer (80 mM PIPES/KOH, pH 6.9, 1 mM MgCl_2_, 1 mM EGTA) containing 20 μM taxol was added to stop the reaction. The sample was subjected to centrifugation at 15,000xg for 20 min at 25°C to collect microtubule seeds in the 200μl prewarmed taxol-BRB80 buffer.

### Single molecule assay by total internal reflection fluorescence microscopy

Imaging was performed using a total internal reflection (TIRF) microscope (Olympus) equipped with an Andor 897 Ultra EMCCD camera (Andor, Belfast, UK) using a 100× TIRF objective (NA 1.49, Olympus). Polymerized microtubules were diluted in the taxol-BRB80 buffer and then flowed into a flow cell and incubated for 10 min at room temperature to adsorb onto the coverslip surface coated with anti-tubulin antibodies (Sigma). Subsequently, polymerized microtubules were protected from laser illumination by flowing the basic reaction mix buffer (RM) (0.08 mg/ml glucose oxidase, 0.032 mg/ml catalase, 0.16 mg/ml casein, 1% β-Mercaptoethanol, 0.001% tween, 1 mM MgCl_2_, 20 μM taxol, 80 mM D-Glucose). Finally, a 50μl final reaction mix consisting of kinesin motors, RM, and 2mM ATP was added to the flow chamber and the time-lapse movies were recorded. Images were taken every 0.1 s with a 0.05 s exposure, and at low laser power to avoid photo-bleaching during processive motor runs.

### Single molecule assay data analysis

In the single molecule motility assay, the position of fluorescent motor spots was manually tracked using the ImageJ software (Fiji, NIH)^58^. In the kymograph, the vertical distance represented the time and the horizontal distance indicated the run length. The ratio between the run length and the time was the velocity. The number of events for each protein sample was plotted for the velocity and run length as a histogram and fitted to a Gaussian distribution. All the statistical analyses were performed using GraphPad Prism (GraphPad Software), and all the data were presented as mean ± SD. All the *P* values were calculated by using a two-tailed unpaired Student *t* test.

### In silico Fisetin docking analysis

The R11Q mutation was introduced into the KIF1A structure (PDB ID: 2ZFI) using PyMOL (Version 2.5, Schrödinger, LLC). The structures of KIF1A(R11Q) and fisetin were minimized and prepared using BioLuminate software (Maestro BioLuminate 4.4, Schrödinger, LLC, New York, NY, 2021). Induced fit docking was then performed between the two molecules, with the sidechain conformation of Tyr67 and Gln70 allowed to be adjusted.

## Supporting information

Example TIRF imaging of GFP-tagged mKIF1A 1-613 (green) sliding along TAMRA labeled taxol-stabilized microtubules (red) immobilized on a coverslip

Example TIRF imaging of 10nM fisetin treated GFP-tagged mKIF1A(R11Q) 1-613 (green) sliding along TAMRA labeled microtubules (red)

Combined TIRF imaging of microtubules gliding along mKIF1A 1-613 of WT (left), R11Q mut (middle) and R11Q mut with 10nM fisetin (right), respectively

## Acknowledgements

This work was supported by the following funding programs: the National Natural Science Foundation of China (grants 32270721, 31991190, 32070706, 32021002, 31730052, 31525015, 31861143042, 31561130153, 31671444, 31671451, 31970180, 31871352, 31900535, 32071191 and 31971160), the National Key R & D Program of China (2017YFA0503501, 2019YFA0508401, and 2017YFA0102900), and the Strategic Priority Research Program of CAS (XDB37020302). The genetic suppressor screens were performed at the 2022 Artificial Evolution Summer School supported by the Tsinghua University and Qinshan Lake Science and Technology City in Hangzhou, China.

---

**Fig. S1.**
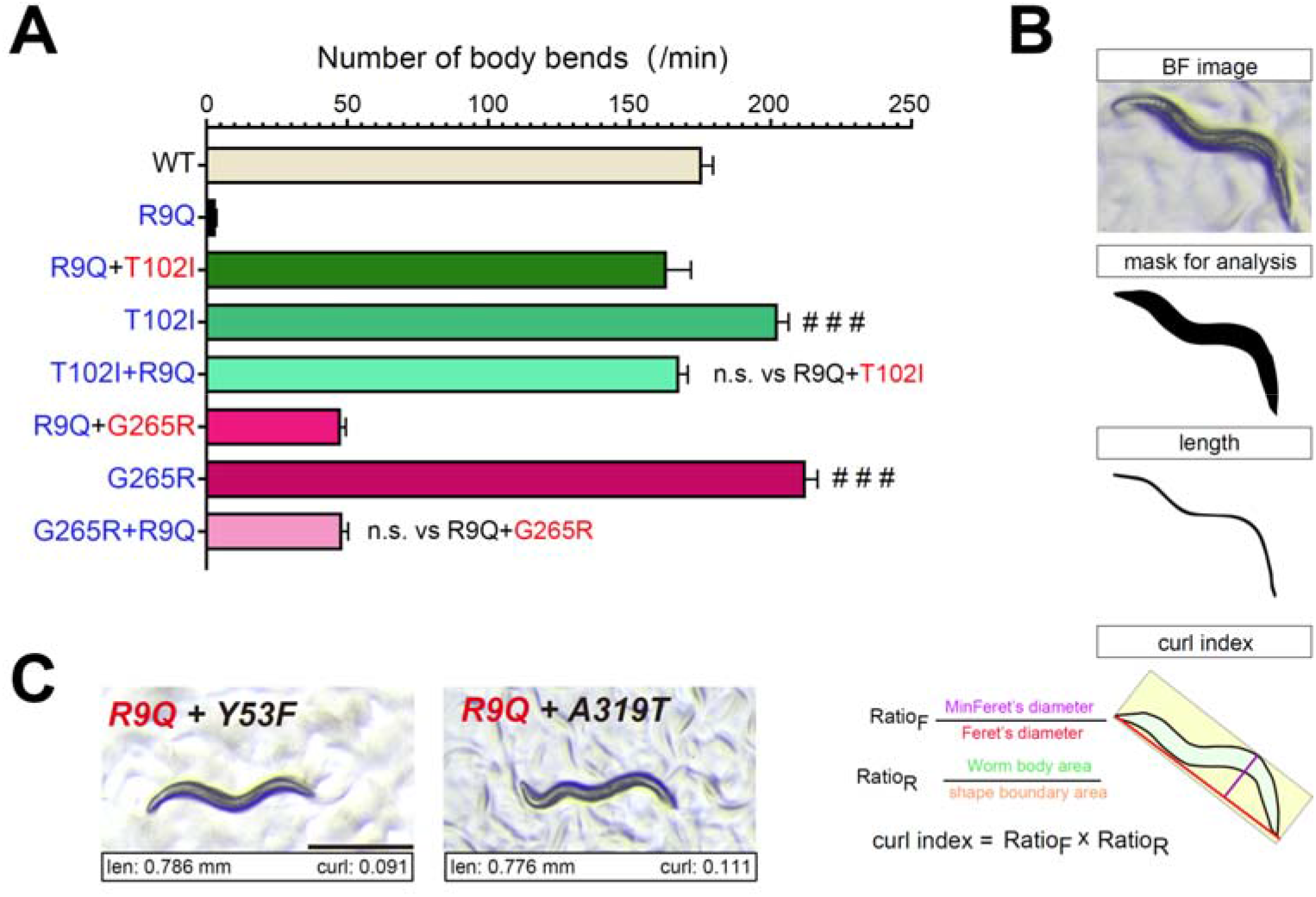
(A) Result of swimming assay conducted with UNC-104 mutant animals bearing amino acid substitutions introduced by CRISPR-Cas9 based genome editing (blue) or identified through the EMS suppressor screen (red). (B) Schematic of workflow used to measure the body length and calculate the body curl index of worms. Tightly entwined animals were excluded from length measurement. (C) Representative bright field images of two homozygous suppressors at the L4 larval stage. The measured body length (len) and calculated curl index (curl) are shown at the bottom. Scale bar, 0.5 mm.

**Fig. S2.**
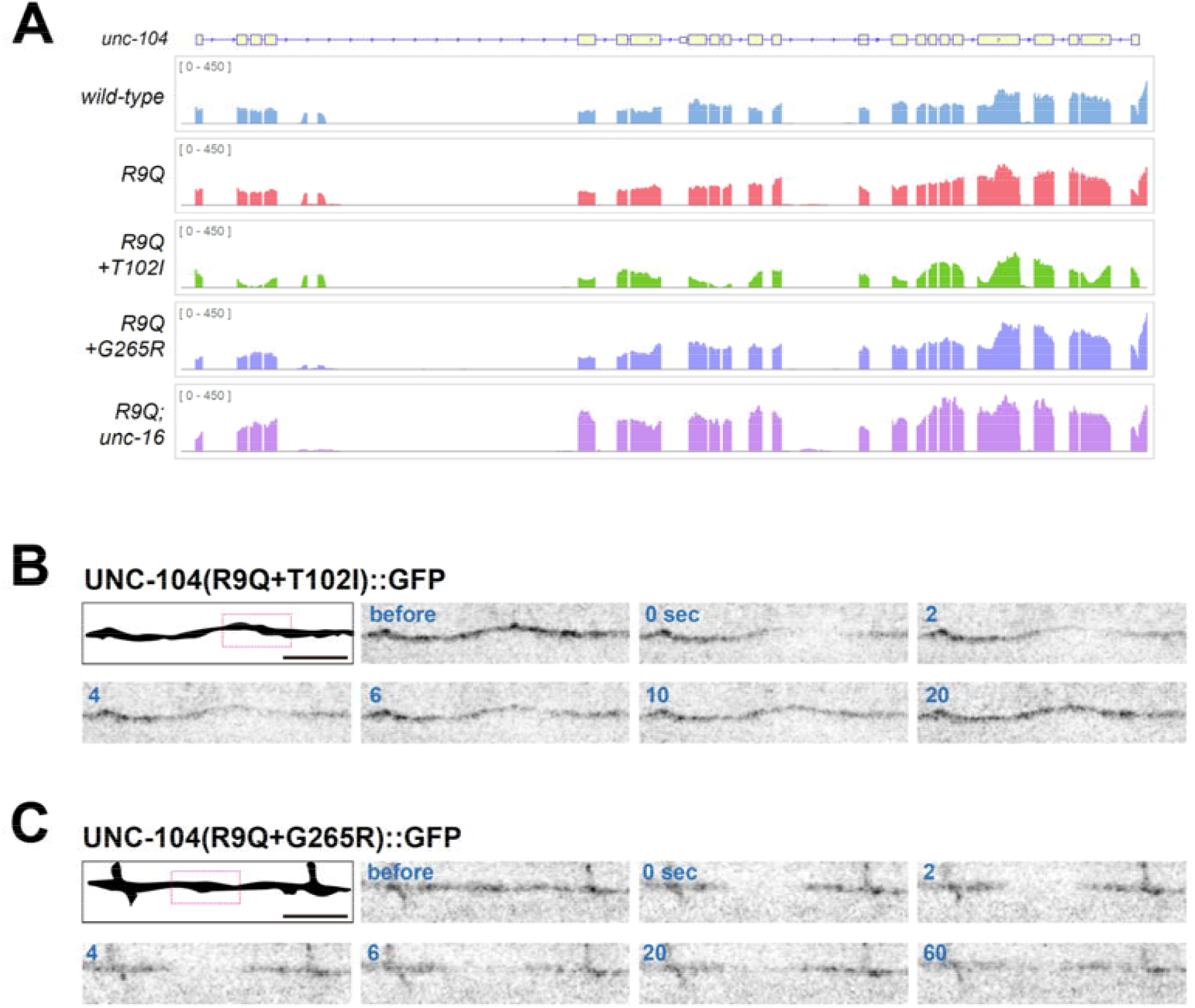
(A) Normalized RNA expression profiles at the *unc-104* locus in different strains as indicated. (B-C) The axons of suppressor UNC-104(R9QT102I):: GFP (B) or UNC-104(R9QG265R):: GFP (C) animals were photobleached within the boxed area of the top left schematic panel, and recovery was recorded for 2 minutes. Images before and after photobleaching (at 0 s) at indicated time points are shown in sequential. Scale bar: 5 _μ_m.

**Fig.S3.**
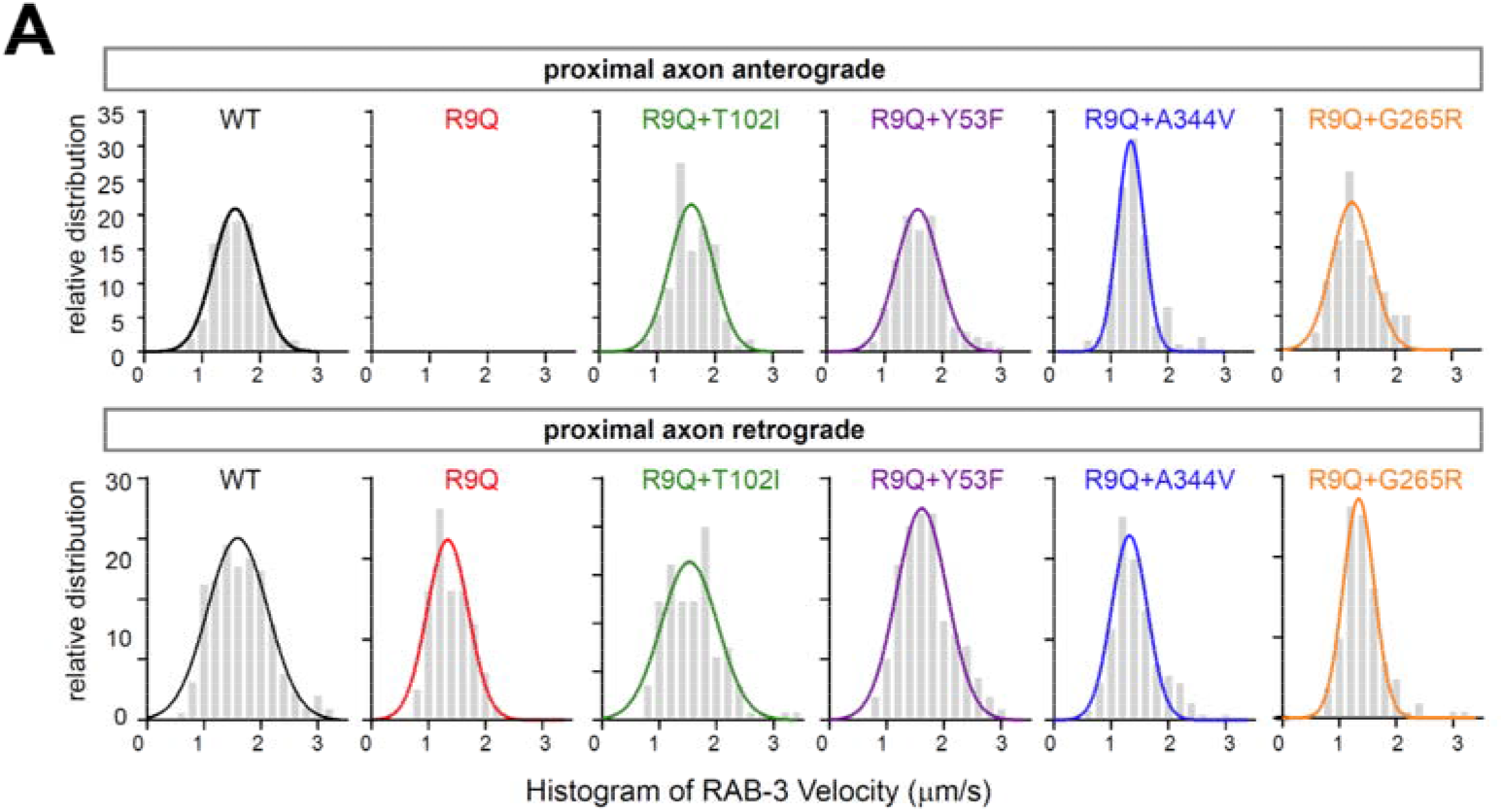
Histograms showing the distribution of anterograde (top) and retrograde (bottom) synaptic vesicle transport velocities in WT, *unc-104(R9Q)* mutant and indicated suppressor animals.

**Fig.S4.**
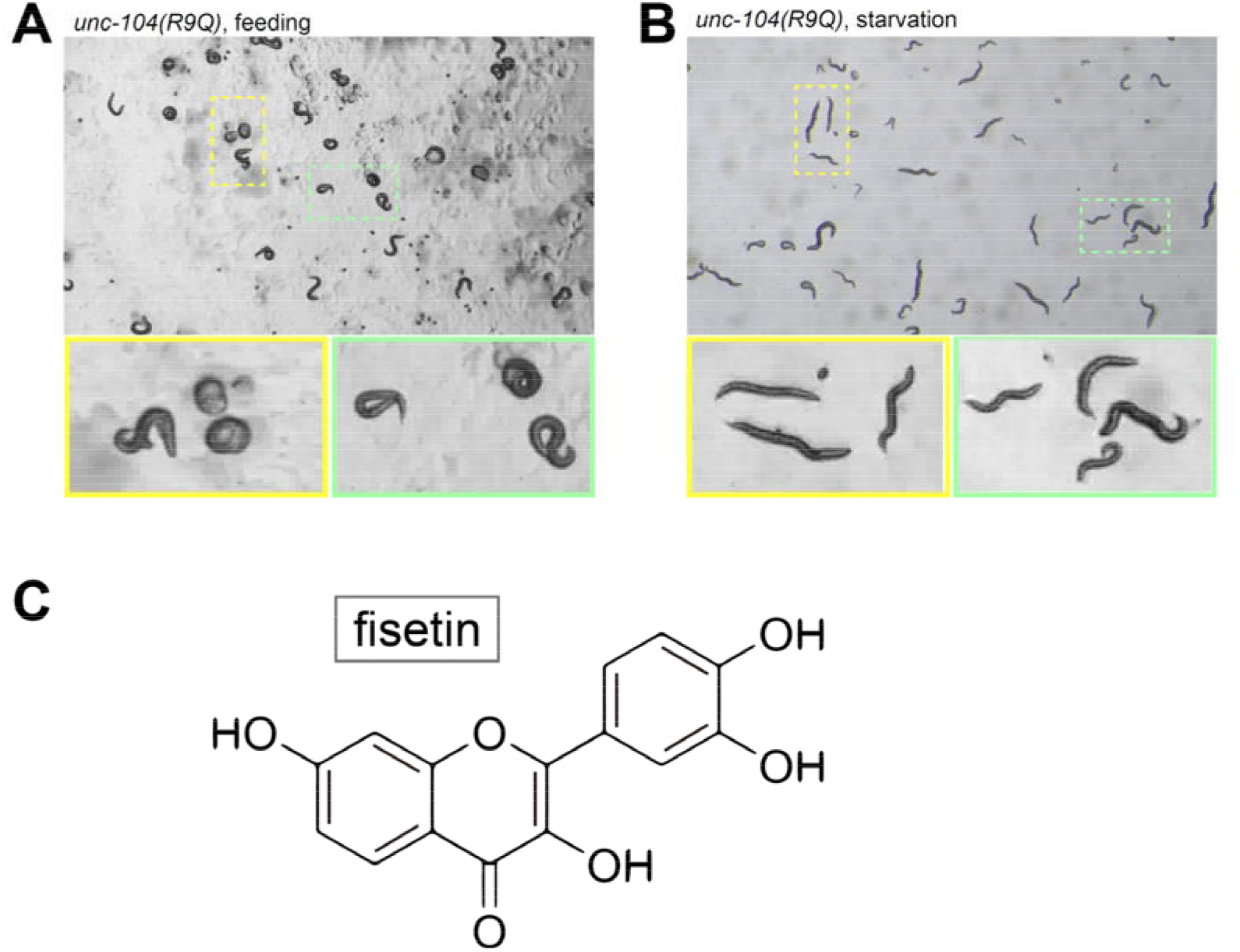
(A-B) Representative bright field images of (A) normal fed and (B) starved *unc-104(R9Q)* mutant animals. The framed areas were enlarged below. (C) Structure of fisetin.

**Fig.S5.**
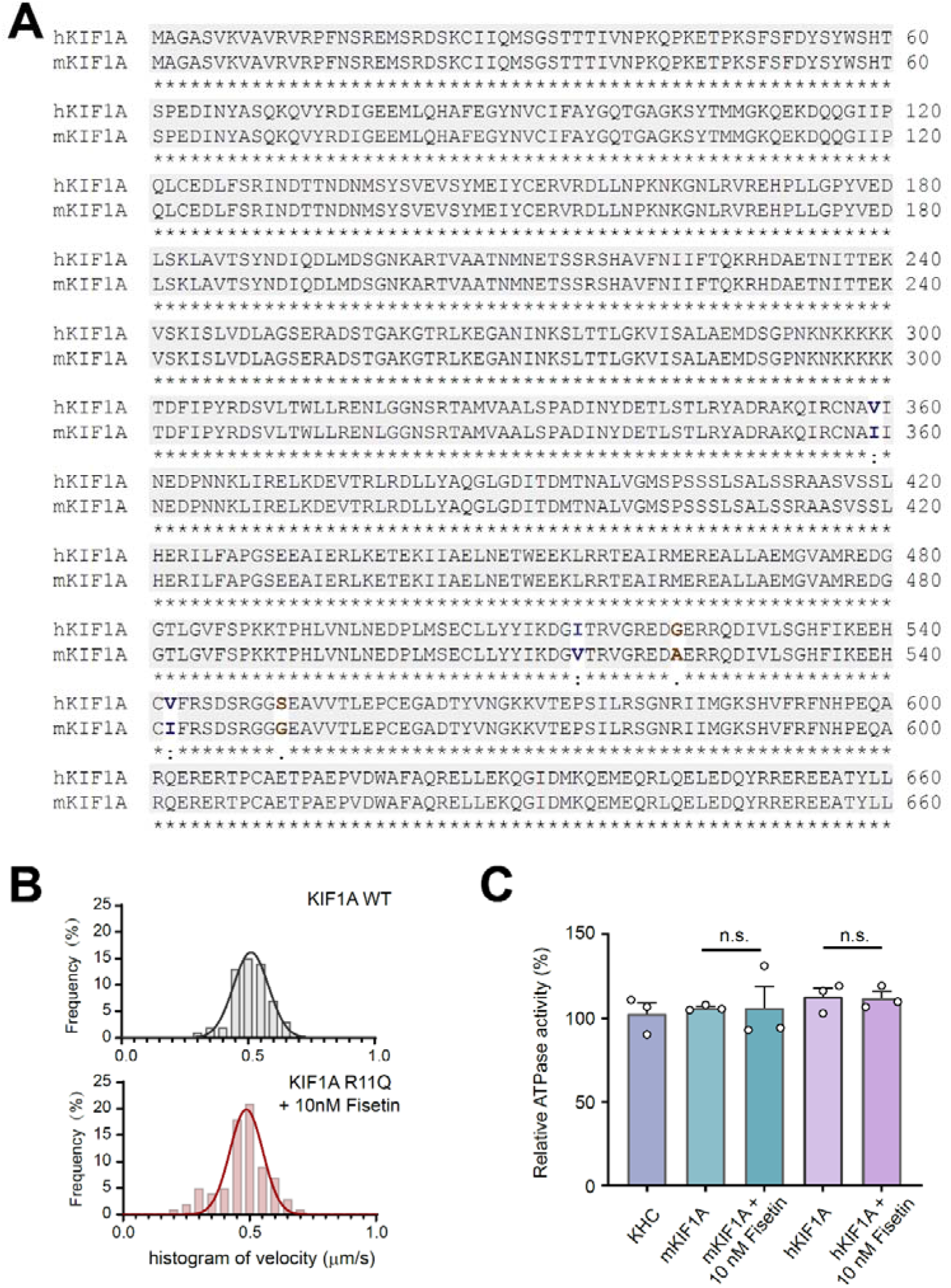
(A) Amino acid sequence alignment of human and mouse KIF1A proteins. Identical amino acids are marked with an asterisk. (B) Qualifications of the microtubule velocity in gliding motility assay for mKIF1A 1-613 (top) and mKIF1A(R11Q) 1-613 motor protein with 10nM fisetin pretreatment (bottom). The number of events is plotted for the velocity as a histogram and fitted to a Gaussian distribution. (C) Microtubule-stimulated ATPase activities of the mouse and human wild-type KIF1A 1-613 protein with or without 10nM fisetin. Data were normalized by the microtubule-stimulated ATPase activity of kinesin-1 heavy chain (KHC) as 100%. Each experiment was repeated three times independently. Each bar represents the mean value ± SD.

## Supplementary Movies

**Video S1.**
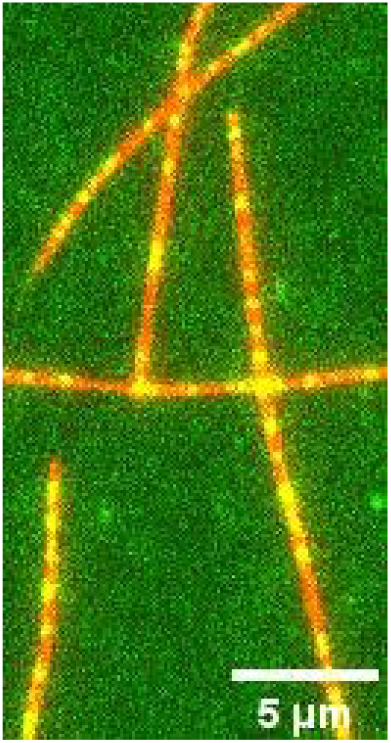
Example TIRF imaging of GFP-tagged mKIF1A 1-613 (green) sliding along TAMRA labeled taxol-stabilized microtubules (red) immobilized on a glass coverslip. Frames were taken every 0.1 seconds. The display rate is 10 frames per second. Details of this event are also shown in Fig. 7C-D.

**Video S2.**
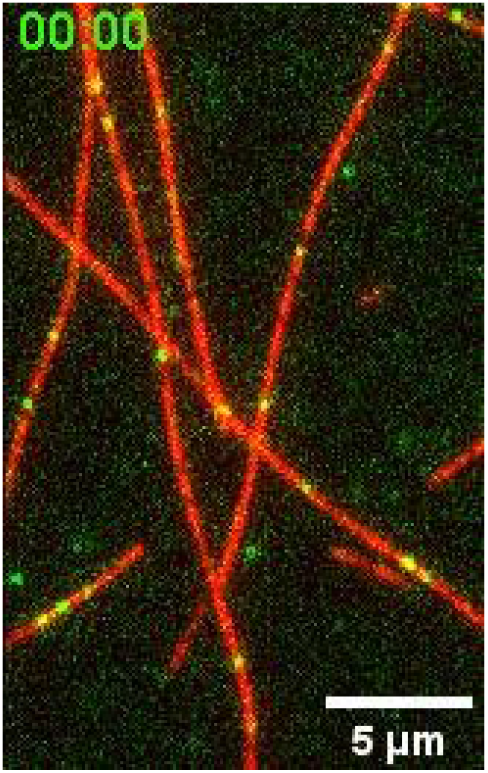
Example TIRF imaging of 10nM fisetin treated GFP-tagged mKIF1A(R11Q) 1-613 (green) sliding along TAMRA labeled taxol-stabilized microtubules (red) immobilized on a glass coverslip. Frames were taken every 0.1 seconds. The display rate is 10 frames per second. Details of this event are also shown in Fig. 7C-D.

**Video S3.**
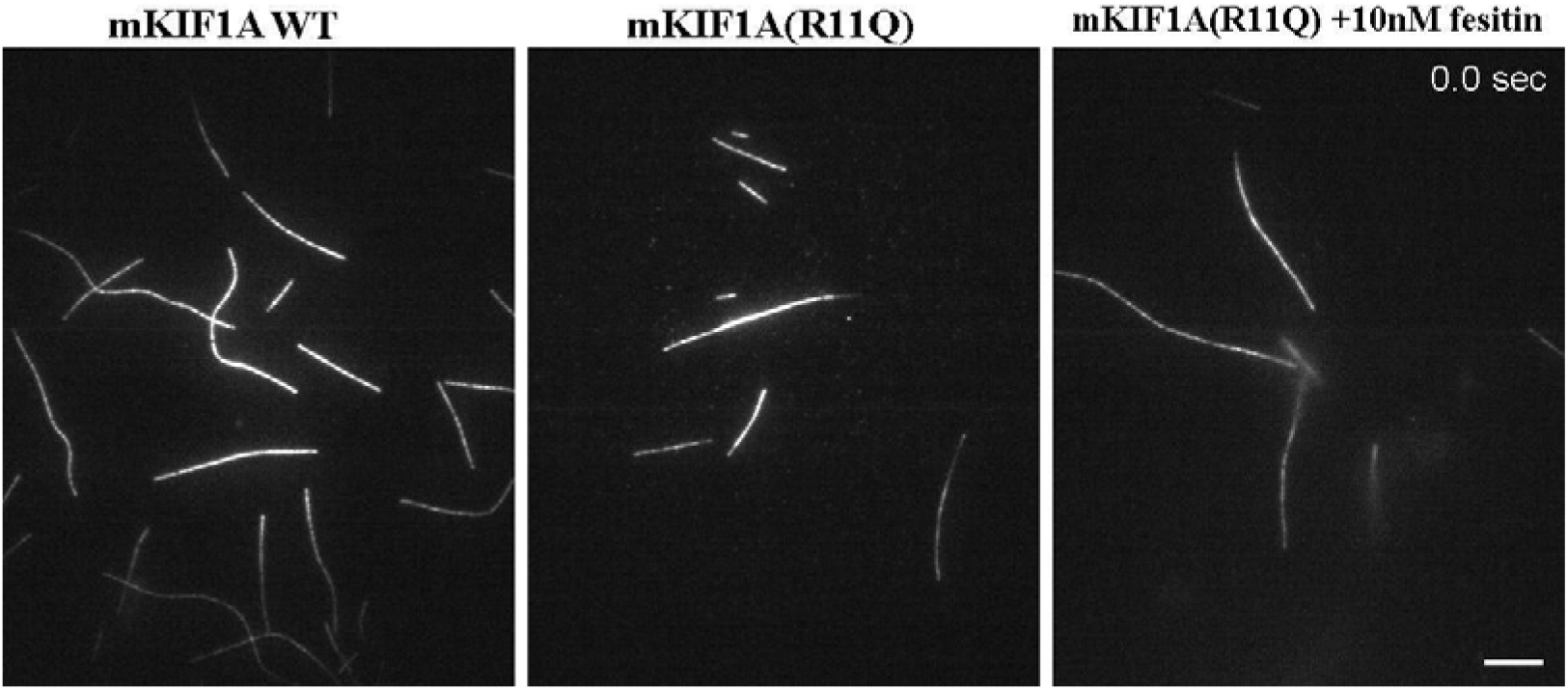
Combined example TIRF imaging of TAMRA labeled taxol-stabilized microtubules gliding along pre-coated GFP-tagged mKIF1A 1-613 protein on a glass coverslip of WT (left panel), R11Q mutant (middle panel) and R11Q mutant with 10nM fisetin (right panel), respectively. Frames were taken every 1 seconds. The display rate is 10 frames per second. Details of this event are also shown in Fig. S5B.

**Table S1.**
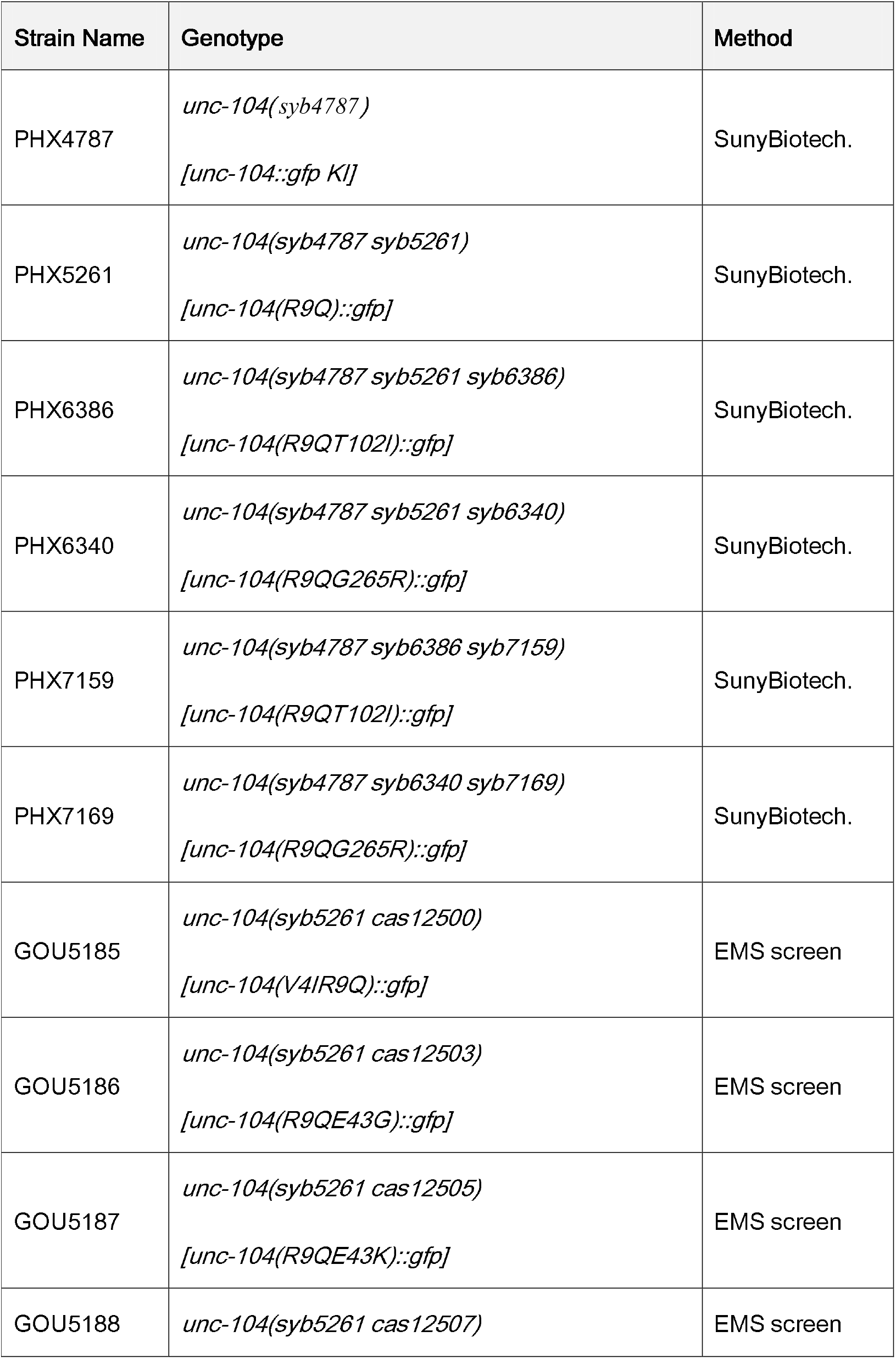
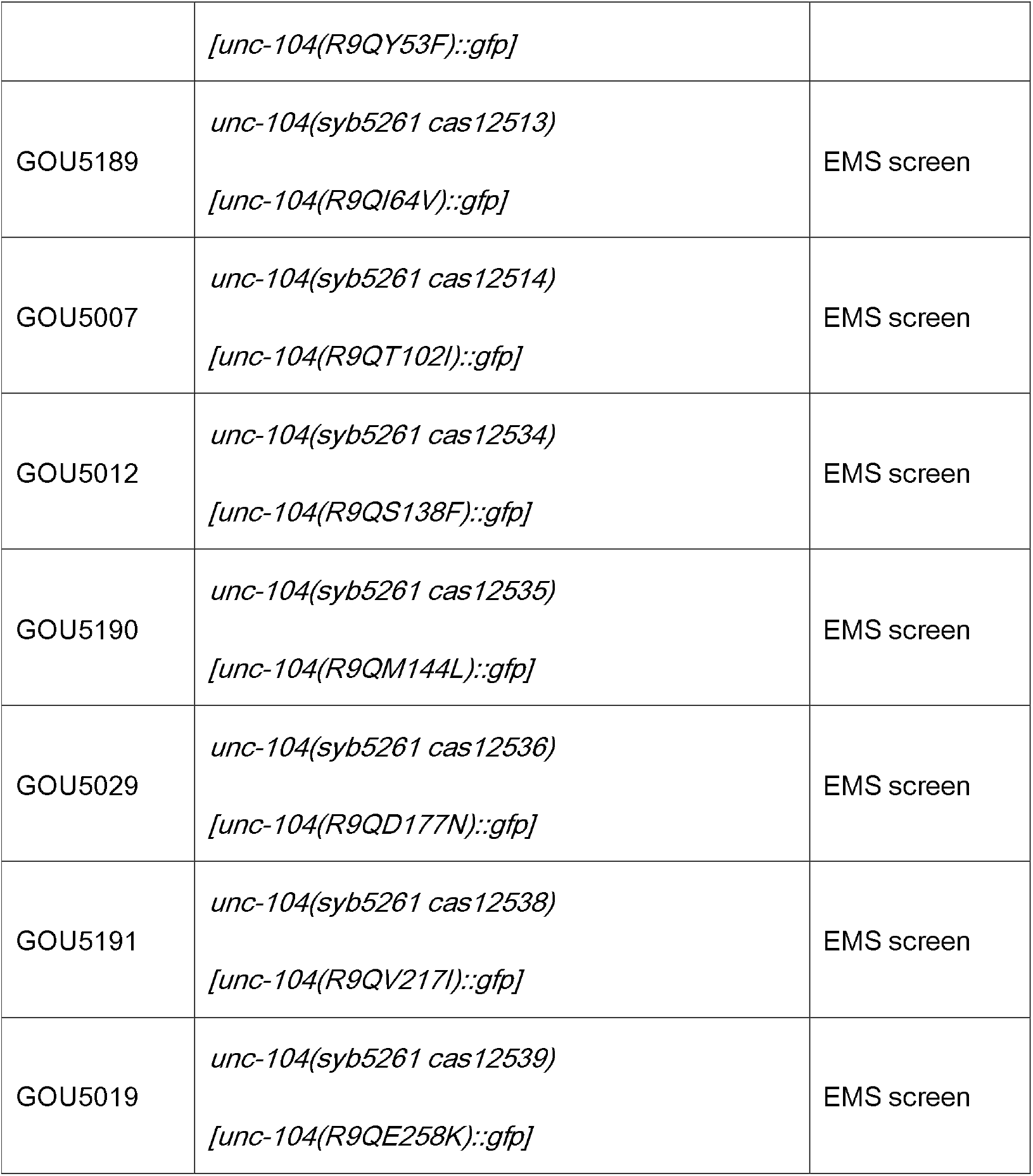
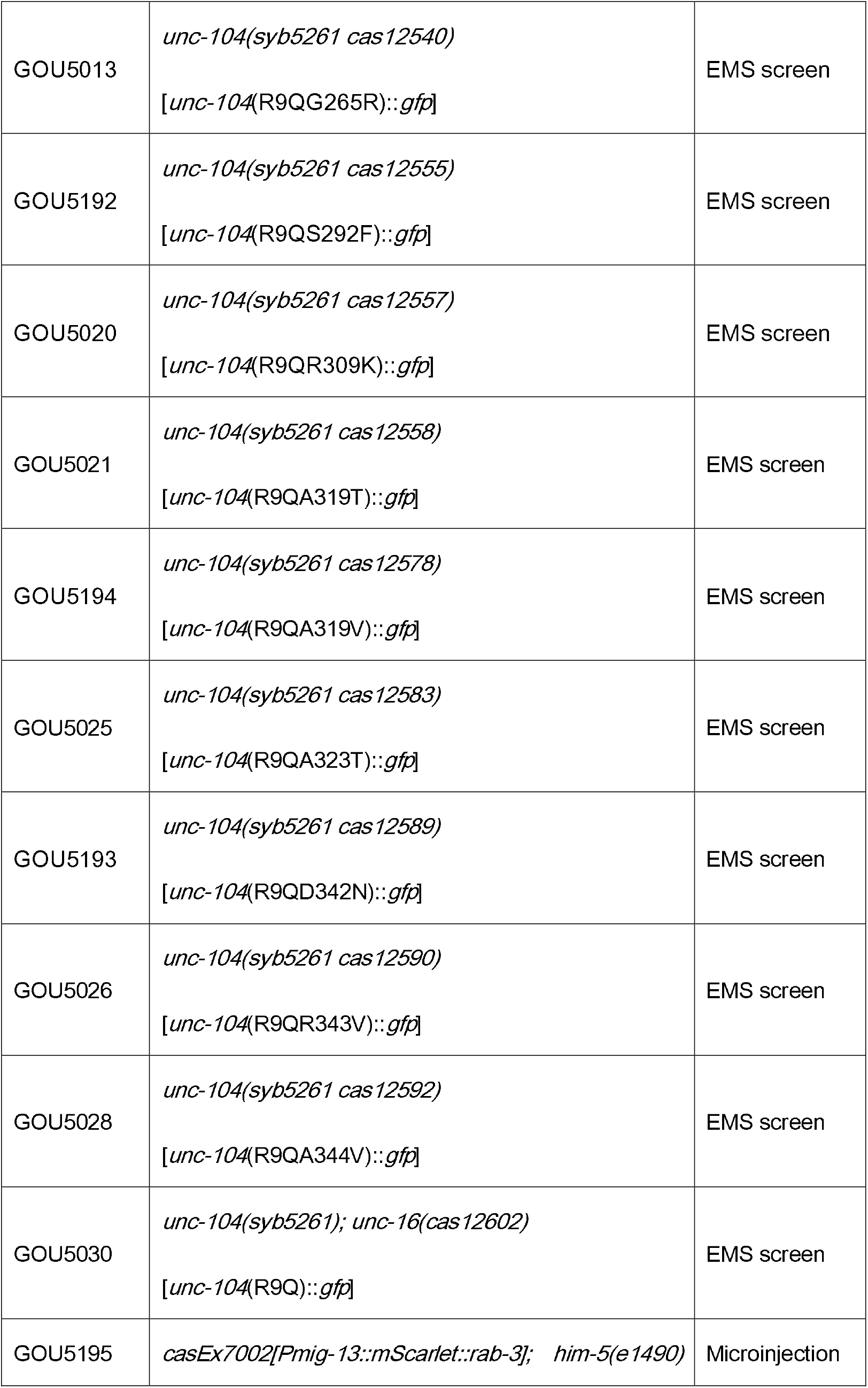
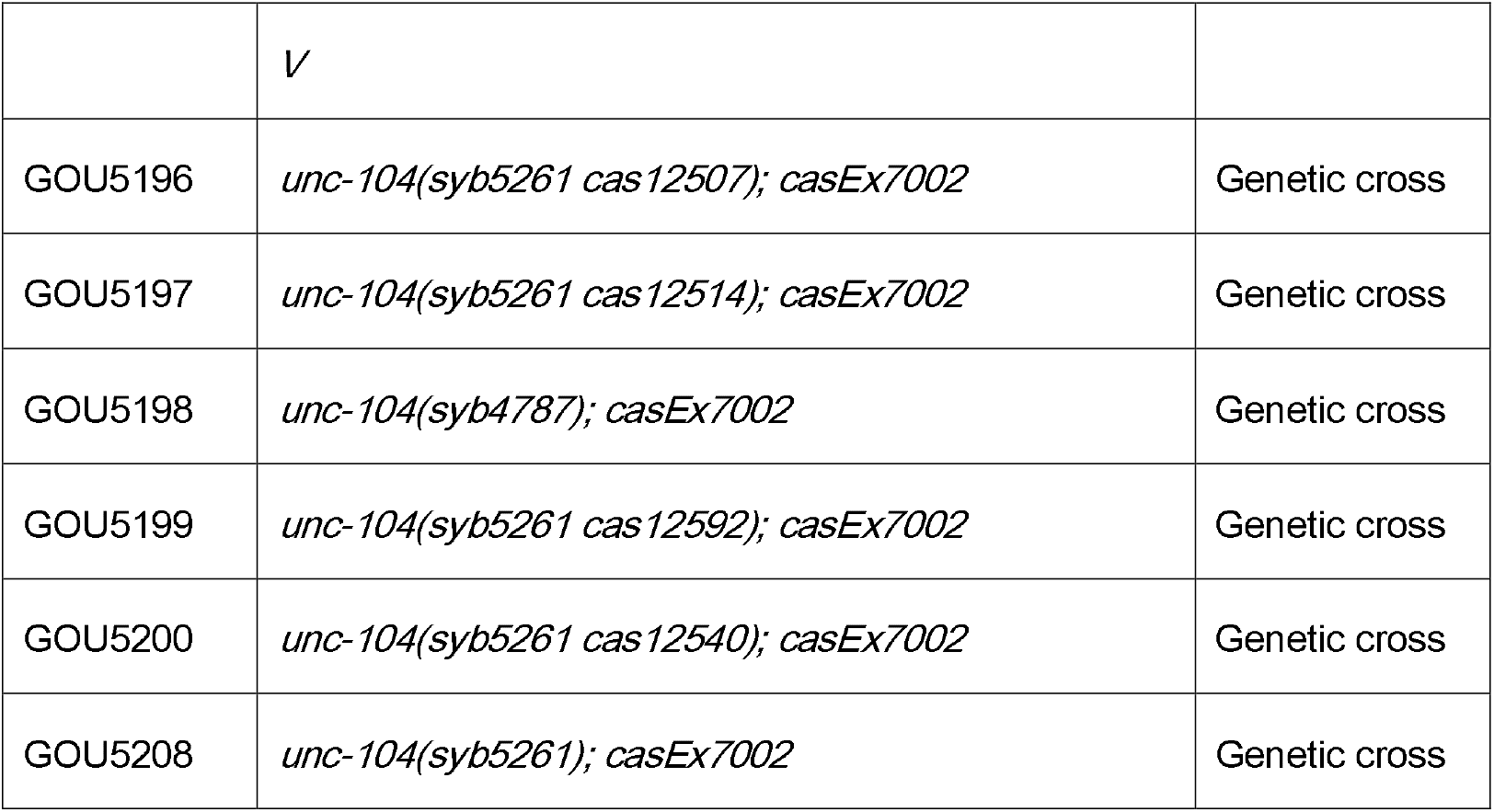
*C. elegans* Strains in this study.

**Table S2.**
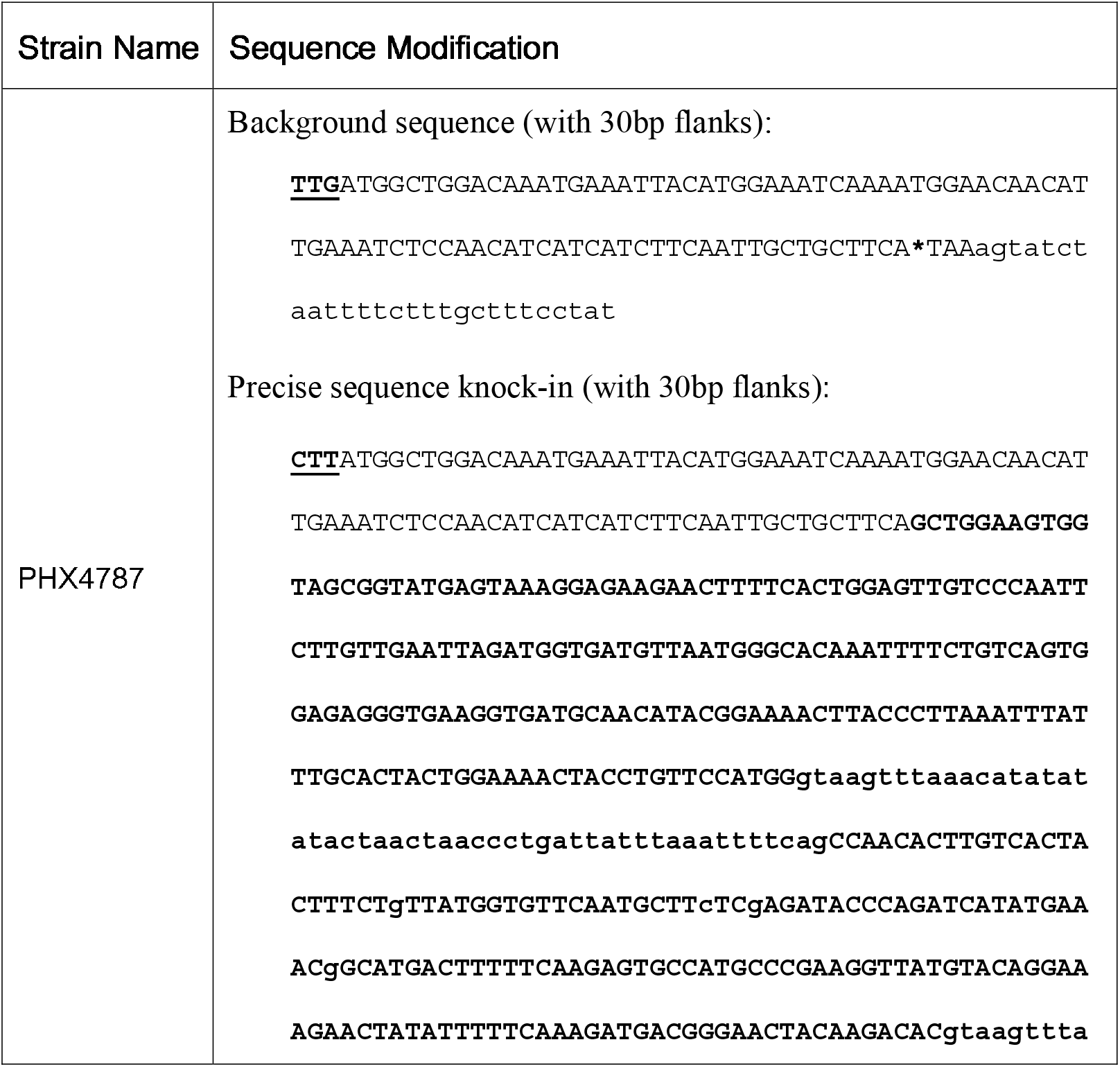
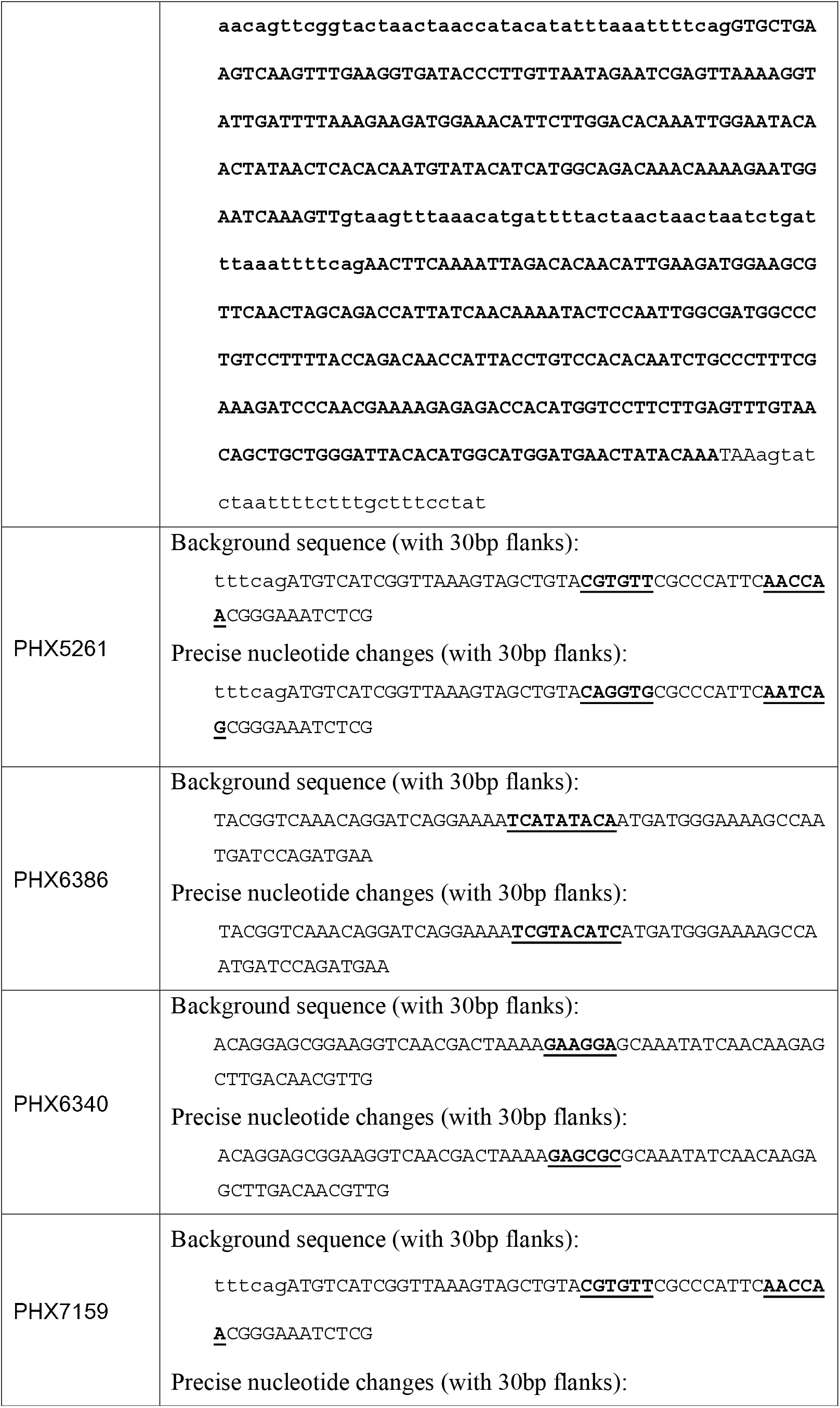
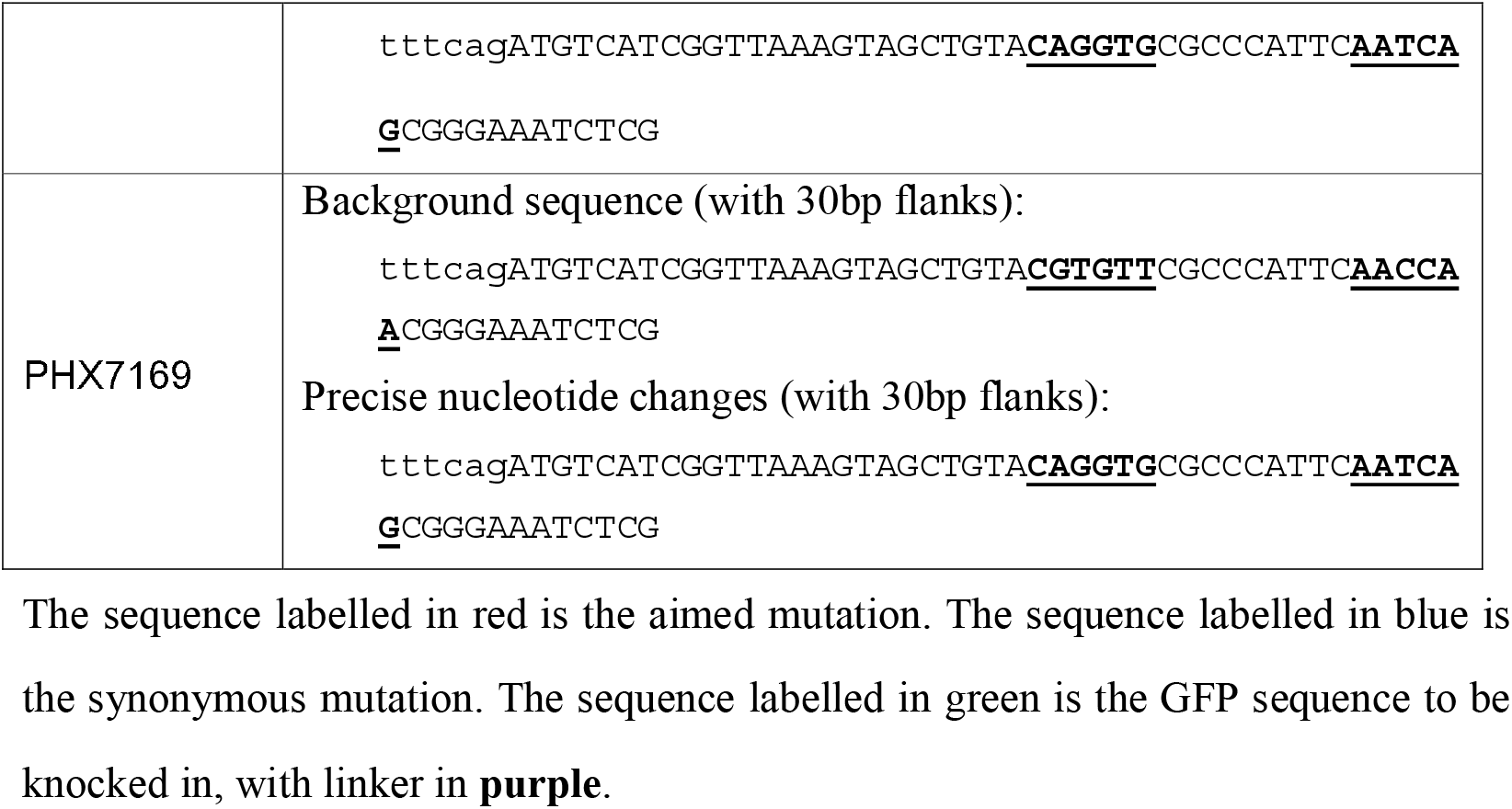
Sequence modification in CRISPR-Cas9 based genome editing.

**Table S3.**
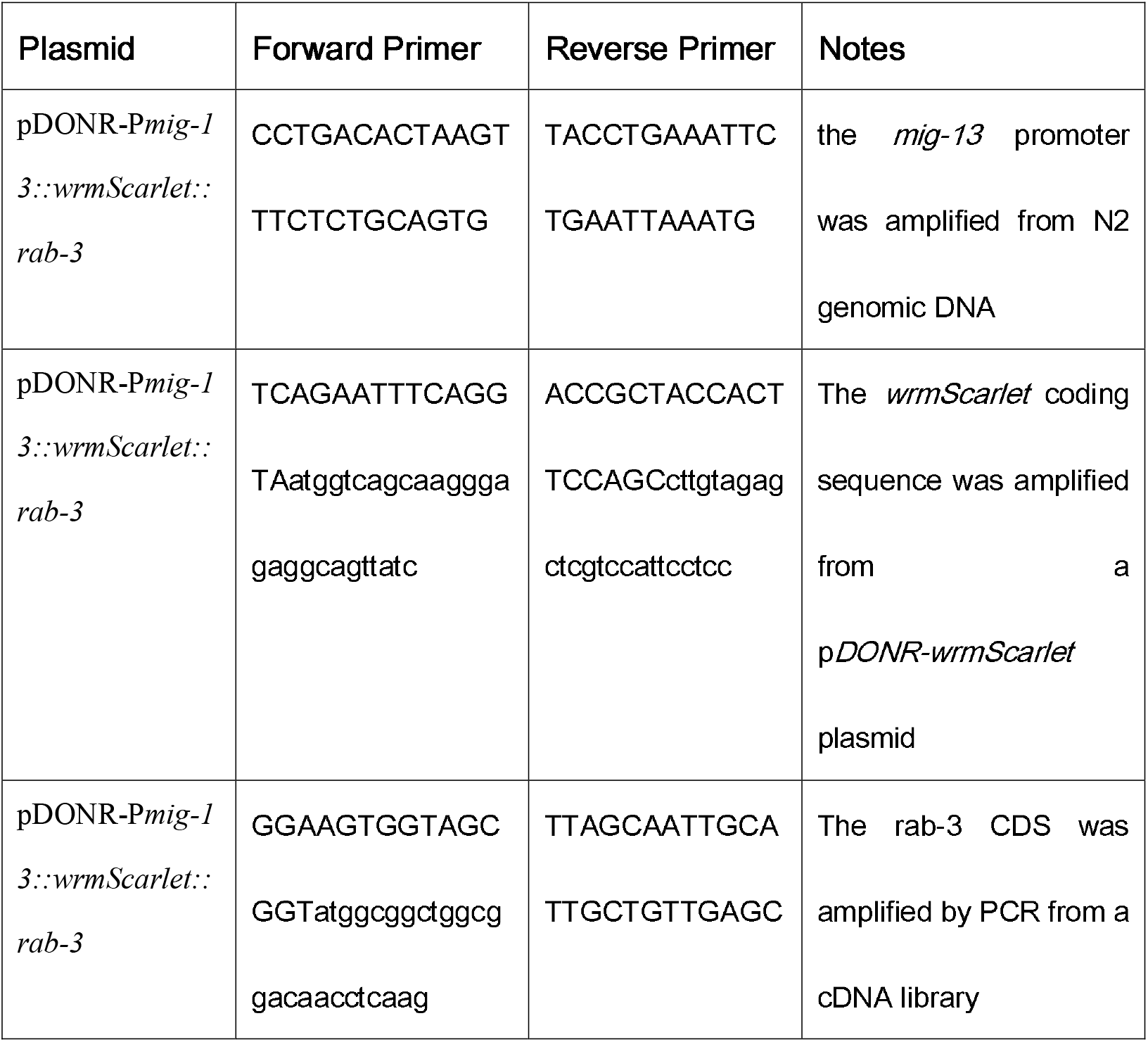
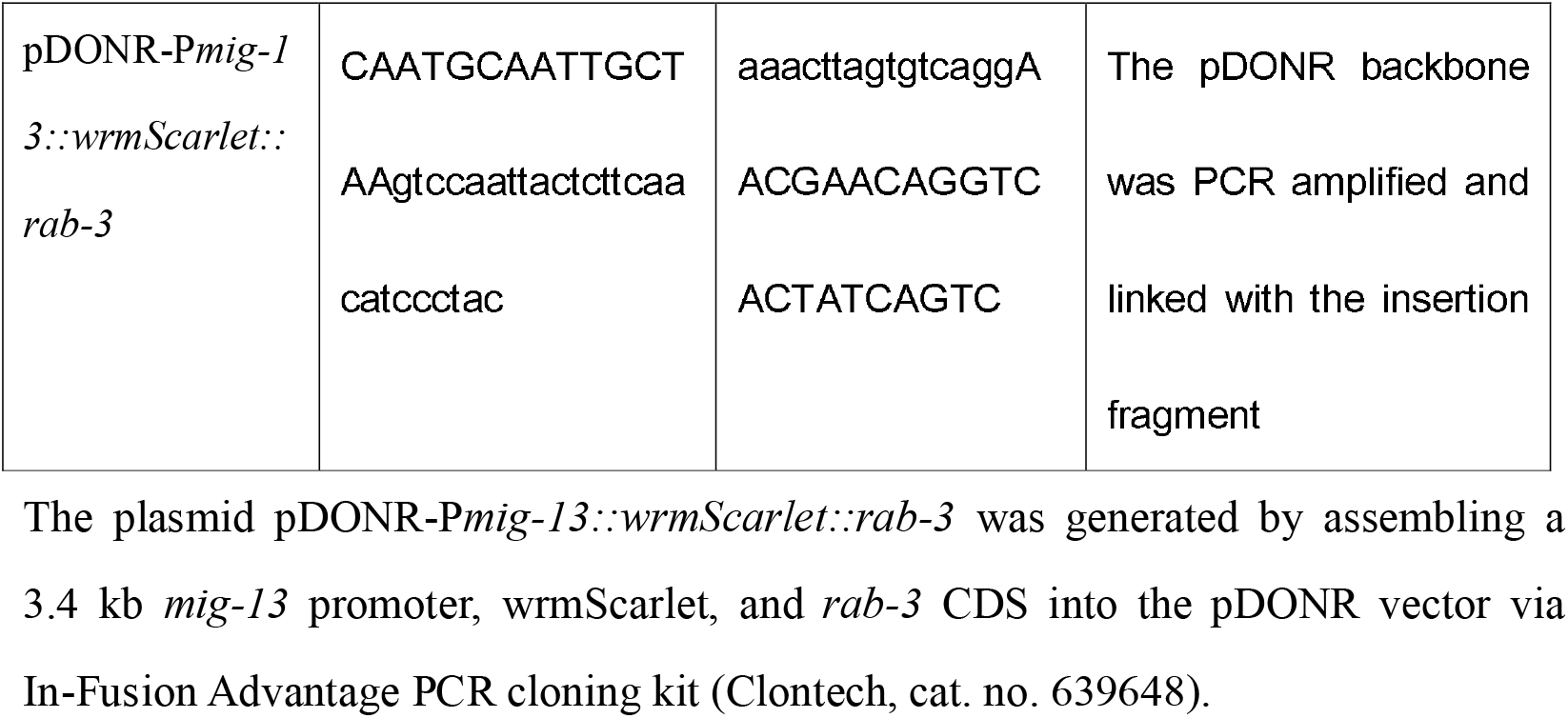
Plasmids and Primers in this study.

